# Pulcherriminic acid biosynthesis and Transport: Insights from a heterologous system in *Saccharomyces cerevisiae*

**DOI:** 10.1101/2025.03.18.643849

**Authors:** Alicia Maciá Valero, Jeroen J. van Wageningen, Alexander J. Foster, Ana Rita Oliveira, Clemens Mayer, Sonja Billerbeck

## Abstract

Pulcherriminic acid is an iron chelator produced by some *Kluyveromyces* and *Metschnikowia* yeasts. Its biosynthesis is encoded by the four–gene *PUL* cluster, where *PUL1* and *PUL2* are the biosynthetic enzymes, *PUL3* mediates the uptake of iron–bound pulcherrimin, and *PUL4* is a putative regulator.

Pulcherriminic acid holds antifungal potential, as the growth of organisms unable to uptake pulcherrimin is inhibited by deficit of essential iron. Thus, a heterologous production system to further characterize and optimize its biosynthesis would be valuable.

Using our in–house yeast collection and genomes available in databases, we cloned *PUL1* and *PUL2* genes from *K. lactis* and one of our wild *Metschnikowia* isolates and built an effective production system in *S. cerevisiae* able to inhibit pathogenic growth. In this context, the *K. lactis* genes yielded faster pulcherriminic acid production than those from the *Metschnikowia* isolate and a combinatorial approach showed *PUL1* to be the production bottleneck.

We further showed that Pul3 is an importer of pulcherrimin, but also mediates the export of pulcherriminic acid and that the growth of pathogens like *Candidozyma auris* and organisms encoding *PUL3* in their genome, previously called “cheaters”, is inhibited by pulcherriminic acid, highlighting its potential as an antimicrobial agent.

## Introduction

Fungal pathogens pose a global threat to human health, food security and agriculture [1], [2] and resistance to all different classes of antifungals has been reported [2]. As such, there is a clear need for novel antimicrobials.

Yeast naturally produce a diverse repertoire of antimicrobial molecules such as killer toxins, hydrolytic enzymes, iron–chelating metabolites, biosurfactants and volatile organic compounds [3], but critical research is lacking to understand these molecules on a genetic and mechanistic level.

In the bacterial world, iron–chelating compounds – commonly referred to as siderophores – recently started gaining attention as potential antimicrobials [4]–[6].

The most studied iron chelators secreted by yeast are pulcherriminic acid (called pulcherrimin in its iron–bound form), produced by *Metschnikowia* and *Kluyveromyces* species (Ascomycota) [7]–[10]; rhodotorulic acid, produced by *Rhodotorula* species but also some filamentous fungi (Basidiomycota) [11]; and fusarinine C (also known as fusigen), produced by the yeast–like fungus *Aureobasidium* (Ascomycota) [12]–[14].

A recent bioprospecting effort undertaken by our laboratory showed that 15% of the isolated yeasts in our collection (107 out of 681 environmental isolates) displayed iron–dependent fungal growth inhibition [15], indicating that this phenotype is rather frequent in nature. Based on the maroon pigmentation of these isolates and their constitutive iron–chelating phenotype in rich media, we hypothesized that most of our isolates would produce pulcherriminic acid or similar iron chelators [16]–[21].

Pulcherriminic acid is a secondary metabolite with a functional hydroxamate group [22]. Although first discovered in the yeast *Metschnikowia pulcherrima* [22], its chemical structure and biosynthesis were first described in the bacteria *Bacillus subtilis* and *Bacillus licheniformis,* where it has been studied for decades [23]–[25]. While the chemical structure of this metabolite is confirmed to be the same in both bacteria and yeast [26], the biosynthetic genes are not related at a sequence level [10]. The gene cluster for pulcherriminic acid production in yeast was first identified and experimentally verified in *Kluyveromyces lactis* in 2018 [10] and subsequently in *Metschnikowia fructicola* (now *M. pulcherrima*) [26], [27]. In both, bacteria and yeast, biosynthesis requires two genes, called *yvmC* or *PUL1* and *cypX* or *PUL2*, respectively. In bacteria, the first biosynthetic step is catalysed by a cyclodipeptide synthase that cyclizes two tRNA–charged leucine molecules into cyclodileucine [28]; the second oxidation step is catalysed by a P450 cytochrome that converts cyclodileucine into pulcherriminic acid. In yeast, *PUL1* is assumed to catalyse the first cyclization step, although it is unclear whether free leucine or tRNA–charged leucine serves as the precursor. The second step is catalysed by *PUL2,* a cytochrome P450 oxidase homolog. The yeast *PUL* gene cluster also encodes for the transporter Pul3 and the putative cluster regulator Pul4 [10]. Pul3 has shown to be essential for uptake of iron–bound pulcherrimin and its presence in the genome of non–producers led to the “cheater” hypothesis [10], where yeasts that do not produce pulcherriminic acid but are able to uptake iron–bound pulcherrimin to counteract the iron–monopolization growth–inhibitory strategy of producers. Such a mechanism could render pulcherriminic acid less useful as an antimicrobial. Freimoser and colleagues previously constructed a heterologous production system based on *PUL1* and *PUL2* from *M. pulcherrima* [29]. However, they reported slower production rates in the heterologous system compared to the natural host.

To address this limitation, we expanded the heterologous production approach by incorporating the *PUL1* and *PUL2* genes from *K. lactis*, the organism where *PUL* genes were first described [10]. This allowed us to construct a fully functional heterologous production system in *S. cerevisiae*, achieving both a reasonable production speed and effective fungal growth inhibition. Our system proved suitable for studying pulcherriminic acid biosynthesis and its potential as an antimicrobial.

Beyond heterologous production, our study aimed to explore the diversity of iron chelator-producing yeasts within our collection. Given the large number of isolates identified, we sought to determine whether these strains exclusively belong to the *Metschnikowia* clade and produce similar enzymes to those previously reported, or if novel producer species and catalytic proteins could be identified. Additionally, we investigated whether a single trait – such as iron chelation – is sufficient to confer antifungal activity against other fungi, including the important human pathogen *Candidozyma auris*.

Finally, we further characterized the role of Pul3, particularly in the context of the “cheater” paradigm [10].

## Materials and Methods

### Materials, equipment and services

All media and buffer components were obtained from BD Bioscience (Franklin Lakes, NJ, USA), Merck KGaA (Darmstadt, Germany) and Thermo Fisher Scientific Inc (Waltham, MA, USA).

Restriction enzymes, ligase and ligase buffer were obtained from New England Biolabs (Ipswich, MA, USA), Phusion Hot Start II DNA Master Mix, GeneArt Gibson Assembly Master Mix and the Pico Green Kit were obtained from Thermo Fisher Scientific Inc (Waltham, MA, USA). Oligos were obtained from Merck KGaA (Darmstadt, Germany), Biolegio (Nijmegen, the Netherlands) and Integrated DNA Technologies IDT (Newark, NJ, USA). Sanger sequencing services were provided by Macrogen Europe (Amsterdam, the Netherlands).

The pulcherriminic standard (CAS 957–86–8) was obtained from MedChemExpress® (South Brunswick, NJ, USA). Sterile tubes and round Petri dishes were obtained from Sarstedt AG & Co (Nümbrecht, Germany). Sterile round–bottom 96–well microtiter plates were obtained from Greiner Bio– One BV (Alphen aan den Rijn, the Netherlands). HTS Transwell®–96 permeable sterile membranes for 96–well plates were obtained from Corning (Corning, NY, USA).

The HPLC system used was a Shimadzu system equipped with an LC–20AD pump, a DGU– 20A3 degasser unit, a SIL–20A autosampler, a SPD–M20A detector, a CBM–20A system controller and a CTO–20A column oven from Shimadzu Corporation (Kyoto, Japan). The column used was an XSelect HSS PFP Column from Waters Corporation (Milford, MA, USA). For LC–MS, we used a UPLC Xevo G2 QTOF system equipped with an ACQUITY UPLC BEH300 C4 column both from Waters Corporation as well.

### Strains and growth conditions

The strains used in this study are listed in **Supplementary Table 1.** *Escherichia coli* was grown at 37°C shaking in LB medium supplemented with chloramphenicol (25 µg/mL) or kanamycin (100 µg/mL) when required.

*Saccharomyces cerevisiae* was cultured at 30°C in Yeast extract Peptone Dextrose medium (YPD: 2.5 g/L yeast extract, 5 g/L peptone, 20 g/L glucose) before transformation. For the selection of transformants, yeast cells were grown at 30°C in optimized synthetic complete (SC Urea) medium unbuffered lacking histidine and uracil (His^-^/Ura^-^) [30]. For the activity screening assays, cells were grown on SC Urea His^-^/Ura^-^ agar buffered to pH 4.0±0.1 with 40 mM Na_2_HPO_4_ and 30 mM citric acid following the 0.2x scheme of the citrate–phosphate buffer system [30].

### Species identification of yeast isolates by Sanger sequencing

The environmental samples were species identified amplifying the Internal Transcribed Spacer 2 (ITS2) region and D1/D2 domain using primers ITS3 and ITS4 [31]; and NL1 and NL4 [32], respectively (**Supplementary Table 2**). The amplicons were cleaned enzymatically using ExoSAP–IT and Sanger sequenced. The sequences were blasted using only type material as a reference and the hits with highest percentage of sequence similarity and query coverage were the species assigned to each yeast sample.

### PUL cluster identification in available genomes

The genome assembly available from the following strains was accessed using the NCBI genome database: *M. rubicola* CBS 15344, *M. pulcherrima* AP47 (previously *M. fructicola*), *M. chrysoperlae* NRRL Y–27615, *M. pulcherrima* KIOM G15050 (previously *M. persimmonesis*) and *C. auris* B11221. Using the online server,genomes were blasted as reference against the genes MPUL0C04990 (*PUL1*), MPUL0C04980 (*PUL2*), MPUL0C04960 (*PUL3*) and MPUL0C04970 (*PUL4*) from *M. pulcherrima* APC1.2 [26] to identify theoretical *PUL* gene clusters for the biosynthesis of pulcherriminic acid.

Gene clusters from *Metschnikowia* species were aligned using Clustal Omega [33] through EMBL Dispatcher [34] to design oligonucleotides for the amplification and sequencing of *PUL* gene clusters in the wild yeast isolates.

All protein sequences were aligned with Clustal Omega via the EMBL Dispatcher as well and visualized with Jalview software [35].

### PUL gene cluster identification and sequencing in wild isolates

A yeast colony PCR was performed using isolates yAMV240, yAMV312, yAMV460, yAMV511, yAMV636 and yAMV642 as template with primers PUL3fw and PUL1rv (**Supplementary Table 2**). The amplicons obtained were cleaned enzymatically using ExoSAP–IT and sent for Sanger sequencing [36] with primers PUL1fw, PUL1midfw, PUL2rv and PUL2midrv.

### Detection of pulcherriminic acid and other secondary metabolites

Wild yeast isolate yMAV511 was inoculated in 50 mL SC Urea medium buffered at pH 4.0±0.1 incubated at 30°C for 24 hours. Then, cells were spun down and the spent media was filter sterilized, freeze dried completely and re–suspended in 1 mL sterile milliQ® water. A standard of pulcherriminic acid was dissolved in DMSO at approximately 2.5 mg/mL concentration.

The 50–fold concentrated yAMV511 spent media and the pulcherriminic acid standard were analysed using a Shimadzu reverse phase high–performance liquid chromatography system. A sample of 10 µL was injected into an XSelect HSS PFP Column (100Å, 5 µm, 4.6 mm x 250 mm) operating at 40°C with a flow rate of 1 mL/min. Separation of the compounds was achieved using 0.1 % (v/v) trifluoracetic acid in water (solvent A) and 0.1 % (v/v) trifluoracetic acid in acetonitrile (solvent B) as eluent. The following gradient was applied: 95% (v/v) solvent A for 5 min, linear gradient to 5% (v/v) solvent A in 15 min, hold at 5 % (v/v) for 3 min; returning to linear gradient to 95 % (v/v) solvent A in 2 min followed by final hold at 95 % (v/v) solvent A for 5 min.

In parallel, samples were analysed by LC–MS on a Waters UPLC Xevo G2 QTOF system. A sample of 2.5 μLwas injected into a C4 column set to 40 ℃. Liquid separation was performed using 0.1 % (v/v) formic acid in water (solvent C) and 0.1 % (v/v) formic acid in acetonitrile (solvent B) as eluent using a flow rate of 0.3 mL/min. The following gradient was applied: 95% (v/v) solvent C for 2 min, linear gradient to 30% (v/v) solvent C in 7 min, linear gradient to 5 % solvent C (v/v) in 1 min, followed by final hold at 5 % (v/v) solvent C for 3 min. Eluted molecules were subsequently ionized by electrospray ionization, positive mode at 3 kV and mass spectra in the 50 to 500 *m/z* range were acquired.

### Molecular cloning and heterologous expression of PUL genes

The *PUL1* and *PUL2* genes from *Metschnikowia* yAMV511 were ordered as synthetic DNA (gBlocks). All CTG codons were modified to AGT (6 and 8 codons, respectively) and codon optimized using the IDT Codon Optimization Tool. Every gene was cloned into pRS413 and pRS416–type plasmids by Golden Gate cloning using the yeast Modular Cloning toolkit (MoClo YTK) [37]. Each gBlock contained overhangs compatible with standard type 3 parts (see **Supplementary Table 3**).

The *PUL1* (KLLA0_C19184g), *PUL2* (KLLA0_C19206g) and *PUL3* (KLLA0_C19250g) sequences from *K. lactis* NRRL Y–1140 [10] were amplified from genomic DNA using specific primers to clone them as Type 3 parts. *PUL2* and *PUL3* were amplified as two fragments to recode BsaI recognition sites present in the original sequence. All primers are available in **Supplementary Table 2**). *PUL1* was cloned into a pRS413–type plasmid and *PUL2* was cloned into a pRS416–type plasmid. Plasmid pPUL2Kl_PUL3Kl containing both *PUL2* and *PUL3* was cloned using the standardized MoClo system for multigene constructs [38].

*PUL1* genes were cloned with TDH3 promoter (pYTK009), ENO terminator (pYTK051) and a *HIS3* marker (pYTK076). *PUL2* genes were cloned with CCW12 promoter (pYTK010), SSA1 terminator (pYTK052) and a *URA3* selection marker (pYTK074). *PUL3* from *K. lactis* was cloned with TEF2 promoter (pYTK014) and ADH1 terminator (pYTK053). All genes were cloned in a low–copy plasmid (CEN6/ARS4 – pYTK081) and in a high–copy plasmid (2 micron – pYTK082) separately, expect for plasmid pPUL2Kl_PUL3Kl, only cloned in a high–copy plasmid. All plasmids used in this study are listed in **Supplementary Table 4**.

All constructs were transformed individually in *E. coli* using standard chemically competent cells and calcium chloride heat–shock procedures and verified by Sanger sequencing.

The constructs successfully cloned were co–transformed in parallel in *S. cerevisiae* and *S. cerevisiae ΔPUL3.* In short, cells were incubated at 30°C in YPD broth and used for the Lithium Acetate PEG transformation protocol [39]. Transformants were grown in selected SC Urea His^-^ /Ura^-^ agar [30] at 30°C. The strains obtained are listed in **Supplementary Table 1**.

### Cell pigmentation and growth inhibition assays

Yeast strains were incubated overnight on 3 mL SC Urea His^-^/Ura^-^ broth pH 4.0±0.1 [30]. For pigmentation assays, the overnight cultures were normalized to 50 final OD_600_ units and 10 µL were spotted on SC Urea His^-^/Ura^-^ pH 4.0±0.1 agar plates with and without 20 µg/mL FeCl_3_ supplementation at 30°C for 24 hours, unless otherwise specified. For growth inhibition assays, the same procedure was followed but with plates containing target strains *S. cerevisiae, S. cerevisiae ΔPUL3* or *C. auris* seeded in agar at 0.01 final OD_600_ units. The plates for growth inhibition were incubated for 48 hours before pictures were taken. Both cell pigmentation and growth inhibition assays were performed in triplicates.

### Protein structure predictions and alignment

Protein structures were predicted using AlphaFold2 [40]. The amino acid sequences used as input were those obtained from Sanger sequencing for Pul1 and Pul2 from *M. pulcherrima* and entries CAH01907.1 and CAH01908.1 for Pul1 and Pul2 from *K. lactis,* respectively. The AlphaFold system used sequence databases to construct a three–dimensional model that was later visualized and aligned using PyMol. The RMSD value was calculated using PyMol as well.

## Results and discussion

### All 30 tested environmental yeast isolates with iron–dependent inhibitory activity belong to the genus *Metschnikowia*

In previous bioprospecting efforts, we isolated 107 yeasts (out of a collection of 681 isolates) that showed iron–dependent inhibitory activity. Based on their maroon pigmentation in the presence of iron and the constitutive character of this phenotype, we suspected them to produce pulcherriminic acid or a similar iron chelator. For taxonomic classification at a species level, we hazardously chose 30 isolates of those 107 for this study, nearly 25% of the iron–dependent inhibitory yeasts. We aimed to identify novel pulcherriminic acid–producing species besides the *Metschnikowia* and *Kluyveromyces* species previously reported [7]–[10].

We first amplified their ITS2 region and used a 97% identity threshold to assign the amplicon obtained to a species and 95% identity threshold to assign it to a genus. All 30 yeast isolates were identified as *Metschnikowia* at a genus level and most were linked to the species *M. pulcherrima* **(Supplementary Table 5)**. The species *M. chrysoperlae, M. rubicola* and *M. pimensis* were occasionally retrieved as well with a slightly lower identity percentage. Given that the *Metschnikowia* genus is known to contain high intra–species diversity in the ITS regions [41], [42], we further amplified the ITS1 region and D1/D2 domain of 10 from those 30 isolates in an attempt to increase taxonomic resolution. However, ITS1 amplicons either showed poor sequencing quality, ambiguity or turned out unreadable as experienced with species from this genus before [43]. The amplification of D1/D2 domains did not improve species resolution either, as it yielded the same species as the ITS2 amplicons **(Supplementary Table 5)**. Consequently, we did not amplify the D1/D2 domains of the 20 remaining isolates and considered only their ITS2 sequencing results for species identification.

As such, 27 out of 30 isolates were assigned to the *M. pulcherrima* clade while isolates yAMV99, yAMV204 and yAMV511 were classified at a genus level as *Metschnikowia* sp. Full genome sequencing would be required to clarify their species level.

Thus, we did not identify novel species producing pulcherriminic acid at this point. However, our results suggest that potentially most of the 107 iron–chelating yeast in our environmentally– sourced collection, if not all of them, belong to the *Metschnikowia* genus. This means that this genus appears to be quite abundant in comparison to other iron–chelating yeast species when studying the iron–dependent growth inhibition phenotype among environmental yeast.

### The architecture of the *PUL* gene cluster and the sequence of its enzymes are conserved within *Metschnikowia* species

We characterized the architecture of the *PUL* gene cluster of six isolates (yAMV240, yAMV312, yAMV460, yAMV511, yAMV636 and yAMV642), assessing the sequence conservation of Pul1 and Pul2 proteins with the goal of assessing if *PUL* clusters were all similar to the ones reported for other *M. pulcherrima* species [26], [27] and cloning *PUL* genes for heterologous pulcherriminic acid production in *S. cerevisiae*.

With the aim of developing a simple PCR assay to analyse the *PUL* gene cluster architecture and perform Sanger sequencing of *PUL1* and *PUL2,* we first compared the genetic organization and gene directionality of the *PUL* clusters from selected genome–available species reported to encode the full four–gene *PUL* cluster (three *M. pulcherrima* strains, *K. lactis* and *C. auris* [10], [26], [44]) and two other *Metschnikowia* (*M. chrysoperlae* and *M. rubicola*) species in which we identified a putative *PUL* cluster (**Figures 1A and 1B**). The multiple sequence alignment showed a conserved *PUL3*–*PUL4*–*PUL2*–*PUL1* architecture for *Metschnikowia* species but differing architectures in *K. lactis* and *C. auris,* both containing a complete *PUL* cluster **(Figure 1A and 1B).**

**Figure 1.**
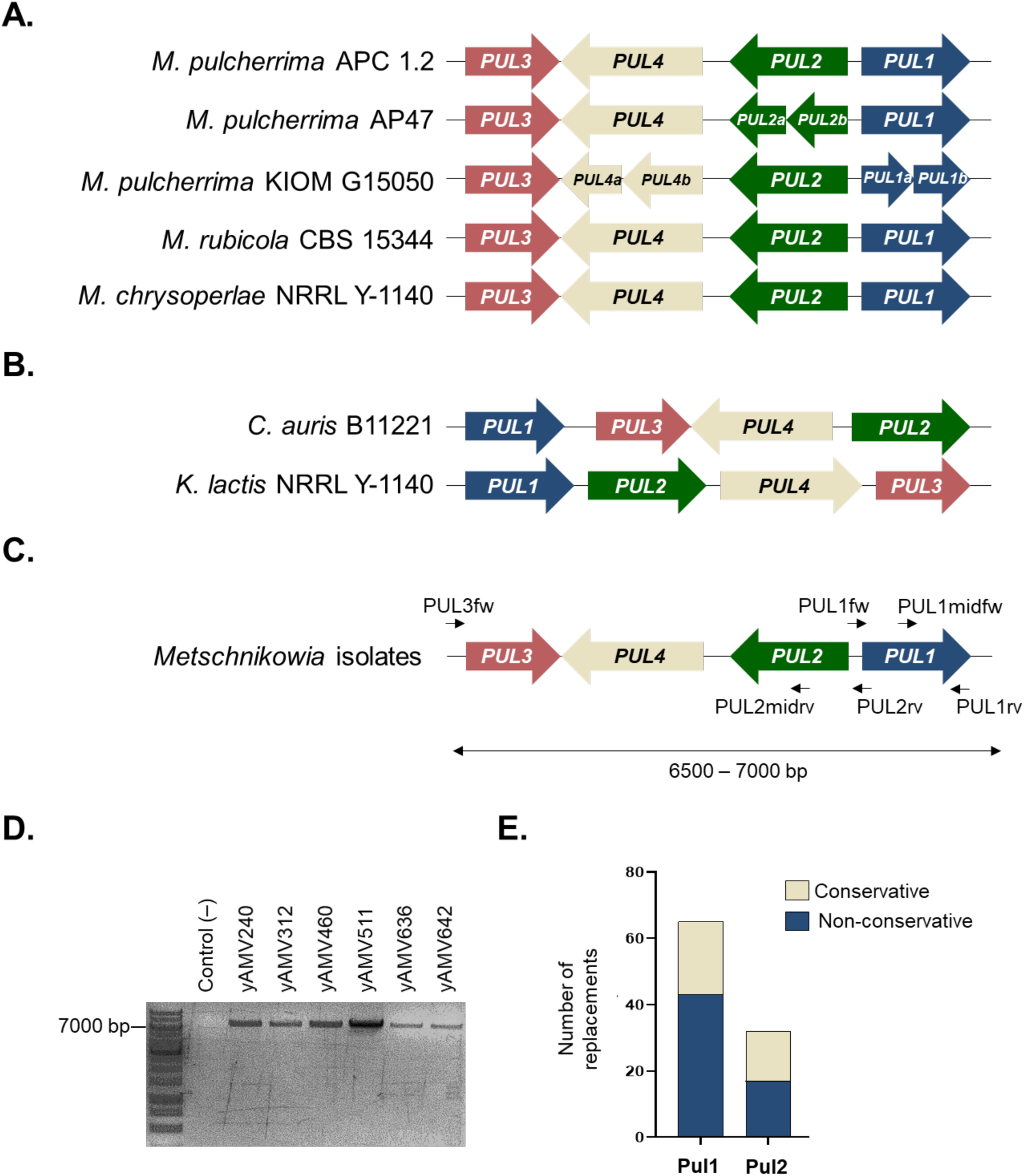
Conservation of the *PUL* gene cluster in architecture and primary amino acid sequence of Pul1 and Pul2 within pulcherriminic acid producers. **A.** The *PUL* cluster architecture within genome–available *Metschnikowia* species. **B.** The *PUL* cluster architecture of *C. auris* and *K. lactis.* **C.** Scheme of the primers designed for the amplification of the *PUL* cluster in yeast isolates and sequencing of *PUL1/2*. **D.** Agarose gel with the amplified *PUL* cluster present in yeast isolates using primers PUL3fw and PUL1rv, all bands show a similar size to other *Metschnikowia* species in bank strains. **E.** Number of conservative and non*–*conservative replacements observed after multiple sequence alignment of Pul1 and Pul2 from *Metschnikowia* strains (in **Supplementary Figures 1 and 2**), including our isolates and genome–available strains.

We then used conserved flanking regions to design primers that would amplify the entire *Metschnikowia* spp cluster and primers for amplifying and Sanger sequencing of *PUL1* and *PUL2* (**Figure 1C** and **Supplementary Table 2**). We obtained a PCR product with an expected size of 6500–7000 bp for all six isolates, indicating the *Metschnikowia*–typical cluster size and architecture with clusters being flanked by *PUL3* and *PUL1* (**Figure 1D**). We then amplified and Sanger sequenced *PUL1* and *PUL2*.

We translated the sequences obtained considering that *Metschnikowia* species belong to the CTG–clade and compared Pul1 and Pul2 primary protein sequences across our isolates and with the sequences from deposited *Metschnikowia* genomes (**Supplementary Table 6**, **Supplementary Figures 1 and 2**). Alignments showed high sequence conservation with sequence identity ranging from 88.76 – 99.78% for Pul1 and 96.46 – 99.79% for Pul2, respectively. Specifically, Pul1 proteins showed 65 variable positions, from which 22 were conservative replacements and 43 were non–conservative replacements. Pul2 showed 32 positions with different amino acids, from which 15 showed conservative replacements and other 17 positions presented non–conservative replacements **(Figure 1E**).

Of note, the *PUL1* locus from *M. pulcherrima* KIOM G15050 – one of the deposited genome sequences – and the *PUL2* locus from *M. pulcherrima* AP47 encoded two open reading frames, aligning to N–terminal and C–terminal portions of the corresponding proteins with a middle segment missing (**Supplementary Figures 1 and 2, Supplementary Table 6**). As such, we did not include them for calculating protein sequence conservation.

We then extended the multiple sequence alignment to *K. lactis* and *C. auris* and checked whether, despite the different gene cluster architecture, Pul1 and Pul2 showed sequence conservation with *Metschnikowia* species (**Supplementary Figures 1 and 2, Supplementary Table 6**). For Pul1, *K. lactis* showed 30.23 – 30.91% sequence similarity with *Metschnikowia* spp while *C. auris* showed 41.20 – 42.12% sequence similarity. For Pul2, *K. lactis* showed 50.00 – 51.26% sequence similarity with *Metschnikowia* spp while *C. auris* showed 56.36 – 56.78% sequence similarity. When comparing *K. lactis* and *C. auris* protein sequences, Pul1 and Pul2 showed 30.77 and 45.89% sequence similarity, respectively. In conclusion, Pul2 showed higher conservation than Pul1 across the yeast species compared.

### High–copy heterologous expression of *PUL1* and *PUL2* genes from *Metschnikowia* isolate yAMV511 is necessary and sufficient for the production of pulcherriminic acid in *S. cerevisiae*

As Pul1 and Pul2 were conserved across our isolates, we picked one isolate, yAMV511, to develop a genetic system for heterologous expression in *S. cerevisiae*.

In media supplemented with iron, our isolate yAMV511 showed a similar cellular pigmentation to previously characterized *M. pulcherrima* yeast producing pulcherriminic acid [26] and its Pul1 and Pul2 primary sequences were almost identical to those of *M. pulcherrima* APC1.2 (**Supplementary Figures 1 and 2**). Using a commercial pulcherriminic acid standard, we confirmed that yAMV511 produced the same iron chelator based on retention time via HPLC and mass peaks on LC–MS (**Supplementary Figures 3 and 4**).

Interestingly, similar to a previous report [26], we detected in the spectrum of the spent media from yAMV511 a peak with a mass of 227.18 *m/z* that was previously hypothesized as cyclodileucine, a precursor of pulcherriminic acid. One would expect this intermediate to only be present intracellularly, so it remains to be elucidated if its extracellular presence originates from active export, leakage or lysed cells, after chemical verification using a cyclodileucine standard.

We then cloned *PUL1* and *PUL2* from yAMV511 into two separate CEN/ARS–based low copy vectors (pRS413–type and pRS416–type) under control of the constitutive strong TDH3 and TEF1 promoters using Golden Gate cloning and parts derived from the yeast MoClo toolkit [37], resulting in plasmids pPUL1Mp_lc and pPUL2Mp_lc (**Supplementary Table 4**). Co– transformation of both plasmids resulted in the *S. cerevisiae* BY4741 strain MpMp_lc.

We tested strain MpMp_lc for pulcherriminic acid production with a simple colorimetric assay on iron–supplemented media [15]. On this media, pulcherrimin producers, such as wildtype isolate *M. pulcherrima* yAMV511 show a maroon pigmentation after 24 hours of incubation (**Figure 2A**). This colour originates from the intracellular accumulation of the maroon–coloured iron–bound pulcherrimin complex. This assay provides a simple qualitative measure for the sum of the following processes: Intracellular pulcherriminic acid production, its export to the extracellular space, iron–binding and maroon pulcherrimin complex formation and import back into the cell. Importantly, *S. cerevisiae* BY4741 encodes a copy of *PUL3* in its genome (Pul3 from yAMV511 and *S. cerevisiae* share 51.49% sequence similarity). Therefore, if *PUL3* is functional and expressed in *S. cerevisiae* BY4741, heterologous constitutive expression of Pul1 and Pul2 should be sufficient for this assay to report results based on cellular pigmentation.

**Figure 2.**
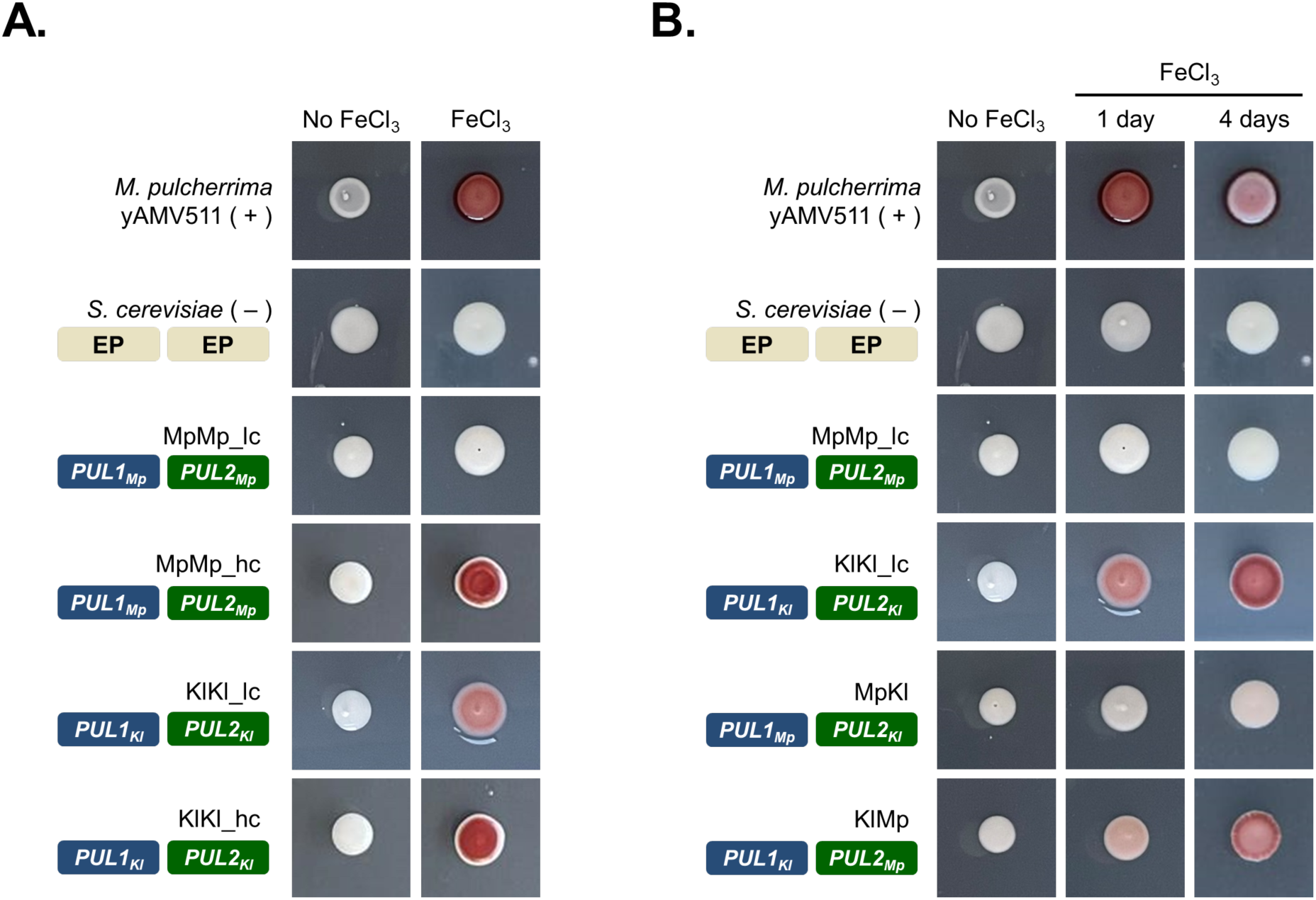
Heterologous pulcherriminic acid production in *S. cerevisiae*. Cell pigmentation assays report pulcherriminic acid production in media with 20 µg/mL FeCl_3_. Wild isolate yAMV511 was used as a positive control (+) and *S. cerevisiae* containing pRS413 and pRS416 empty plasmids (‘EP’) was used as a negative control (–). Pictures are representative of results from assays performed in triplicate. **A.** Heterologous expression of *PUL1/2* from isolate yAMV511 in a low copy (MpMp_lc) and high copy plasmid (MpMp_hc) and *PUL1/2* from *K. lactis* in a low copy (KlKl_lc) and high copy plasmid (KlKl_hc). Cells were incubated for 24 hours at 30°C, **B.** Combinations of *PUL1/2* genes from *Metschnikowia* isolate yAMV511 and *K. lactis.* Cells were incubated for 4 days at 30°C.

For our colorimetric assay, we used isolate yAMV511 as a positive control and *S. cerevisiae* co–transformed with pRS413 and pRS416 empty plasmids as a negative control. When growing spots of *S. cerevisiae* MpMp_lc heterologously expressing *PUL1/2* from *Metschnikowia* isolate yAMV511, we did not see any cell pigmentation after 24h (**Figure 2A**) or even a prolonged incubation up to 14 days.

During the course of this project, Freimoser and colleagues reported the heterologous expression of genome–integrated *M. pulcherrima PUL1* and *PUL2* in *S. cerevisiae* BY4741 [29] showing that, in their system, an incubation of seven to 14 days was necessary for any cell pigmentation to be observed and that gene copy number correlated with the intensity of cell coloration. In their system, 1 to 5 genomic copies of the *PUL1/2* genes were sufficient for the production of pulcherriminic acid; while for our system, this copy number (CEN/ARS, 2 to 5 copies [45]) was insufficient, despite the use of similar promoter and terminator combinations from the MoClo YTK. Of note, their experiments were performed at 22°C [29], growth conditions more suitable for *M. pulcherrima;* while our incubation temperature was 30°C, the standard *S. cerevisiae* temperature. Thus, it is possible that the stability or activity of *Metschnikowia* Pul1 and Pul2 proteins is optimal at a lower temperature.

To test if increased copy number would yield visible pulcherriminic acid production, we built plasmids pPUL1MP_hc and pPUL2Mp_hc cloning *PUL1* and *PUL2* in high–copy 2 micron plasmids (14 to 34 copies[45]) with the same promoters and terminators (**Supplementary Table 4**). Transformation yielded *S. cerevisiae* strain MpMp_hc. This strain showed the characteristic maroon cell coloration after 24 hours of incubation on iron–supplemented media (**Figure 2A**), confirming that a high copy number was required for sufficient production of pulcherriminic acid to be visible in our assay.

In summary, we confirmed that *PUL1* and *PUL2* genes are sufficient for pulcherriminic acid production in *S. cerevisiae* and that the endogenous copy of *S. cerevisiae PUL3* (*YNR062C*) gene is functional and expressed as it was previously suggested [10].

### *PUL1* and *PUL2* genes from *K. lactis* constitute a more efficient genetic system for the production of pulcherriminic acid in *S. cerevisiae* and reveal *PUL1* as a production bottleneck in the system from *Metschnikowia* yAMV511 isolate

In a parallel project, we had built with *PUL1/2* genes from *K. lactis* the plasmids pPUL1Kl_lc and pPUL2Kl_lc into the same low–copy plasmids and the plasmids pPUL1Kl_hc and pPUL2Kl_hc into high–copy plasmids, all of them using the same promoters as above (**Supplementary Table 4**). Co–transformation into *S. cerevisiae* yielded strains designated as KlKl_lc and KlKl_hc, respectively. Strikingly, strain KlKl_lc showed cell pigmentation after only 24 hours of incubation at 30°C (**Figure 2A**), as opposed to strain MpMp_lc that had not shown any pigmentation after 14 days. Moreover, strain KlKl_hc, as expected, yielded a more intense pigmentation than KlKl_lc, similar to MpMp_hc against MpMp_lc, after only 24–hour incubation. In conclusion, *PUL1* and *PUL2* genes from *K. lactis* were heterologously expressed for the first time and resulted in a more efficientagenetic system for the study of *PUL* genes in *S. cerevisiae* when compared to the expression of *PUL* genes from our *Metschnikowia* isolate. The dysfunctional low–copy *PUL1/2* expression system from yAMV511 (strain MpMp_lc) and the functional low–copy *PUL1/2* expression system from *K. lactis* yielded the opportunity to identify a bottleneck in expression by creating a combinatorial expression system.

Thus, we created *S. cerevisiae* strain MpKl by co–transforming plasmids pPUL1Mp_lc and pPUL2Kl_lc, and strain KlMp by co–transforming plasmids pPUL1Kl_lc and pPUL2Mp_lc (**Supplementary Table 4**).

**Figure 2B** shows that, after 24 hours of incubation, strain KlMp displayed some cell pigmentation, developing a stronger coloration after 4 days; while strain MpKl, did not show any pigmentation, indicating that Pul1 from yAMV511 constituted a constraint in the production of pulcherriminic acid in *S. cerevisiae* in this combinatorial setup.

We reported above that the protein sequence of Pul1 is less conserved across some yeast than the sequence of Pul2. In addition, we performed structure predictions of yAMV511 Pul1/2 and *K. lactis* Pul1/2 using AlphaFold2. The predictions yielded structures of Pul1 and Pul2 from yAMV511 with average pLDDT scores of 72.69 and 91.22, respectively, and structures of Pul1 and Pul2 from *K. lactis* had average pLDDT scores of 85.04 and 88.53, respectively. The structure prediction was followed by structural alignments of those predictions of each protein from both organisms. For the two Pul1 structures, the RMSD value was 1.543 Å; for the two Pul2 structures, 0.904 Å, again indicating more variation within Pul1 than Pul2 (predicted structure alignments in **Supplementary Figures 5A and 5B).** Additional in–depth enzymatic studies would be required to investigate whether *Metschnikowia* Pul1 expression levels, folding or catalysis are constraining the pulcherriminic acid production.

### High–copy number–based expression of *PUL1* and *PUL2* from *K. lactis* is necessary and sufficient to replicate the iron–dependent growth inhibition phenotype in *S. cerevisiae*

Next, we tested whether the heterologous production of pulcherriminic acid was sufficient to cause growth inhibition of other yeasts, thus replicating the full phenotype of the iron– monopolizing yeast isolate yAMV511. This is important, as other factors, such as competition of nutrients or production of other secondary metabolites, might contribute to the growth inhibition caused by environmentally isolated strains. The heterologous production also allows to test for the antifungal capacity of pulcherriminic acid in a clean strain background.

This time we used a Halo assay to test for growth–inhibitory activity. As previously described [15], this assay is similar to the above presented but includes a sensitive strain embedded in agar plates (in this case, *S. cerevisiae*) and it is performed with and without iron supplementation. This assay has a dual readout, testing for the production of pulcherriminic acid – including export, iron binding and pulcherrimin uptake – via maroon-colour formation of the producer and growth inhibition, showing as a growth inhibition zone (halo) around the producer.

We used *S. cerevisiae* as a target strain because, despite containing a functional *PUL3* gene and being a potential cheater, because it had shown sensitivity against iron–chelator producing yeast isolates in a previous study [15]; which retrospectively was surprising to us because we would expect it to be a non-sensitive cheater.

Pulcherriminic acid producing *S. cerevisiae* BY4741 (low– and high–copy production strains KlKl_lc and KlKl_hc) were spotted on assay plates with embedded sensitive strain in the presence or absence of iron. Spots of the wild isolate yAMV511 were used as a positive control and spots of *S. cerevisiae* BY4741 transformed with empty plasmids as a negative control. A growth inhibition halo was only expected to appear on plates with trace amounts of iron, where iron–binding by pulcherriminic acid depletes the indicator strain of this essential nutrient, while pulcherrimin transport back into cells supports the producer. We would expect this inhibition to be alleviated in the presence of high amounts of iron. Both phenotypes could be successfully shown with the positive control (**Figure 3A**). The high–copy–based expression system led to the formation of a growth inhibition halo (KlKl_hc) and growth inhibition disappeared in the presence of high iron, as expected. For the low–copy heterologous expression strain KlKl_lc, no halo was observed indicating that this strain did not produce enough pulcherriminic acid to inhibit the growth of *S. cerevisiae*.

**Figure 3.**
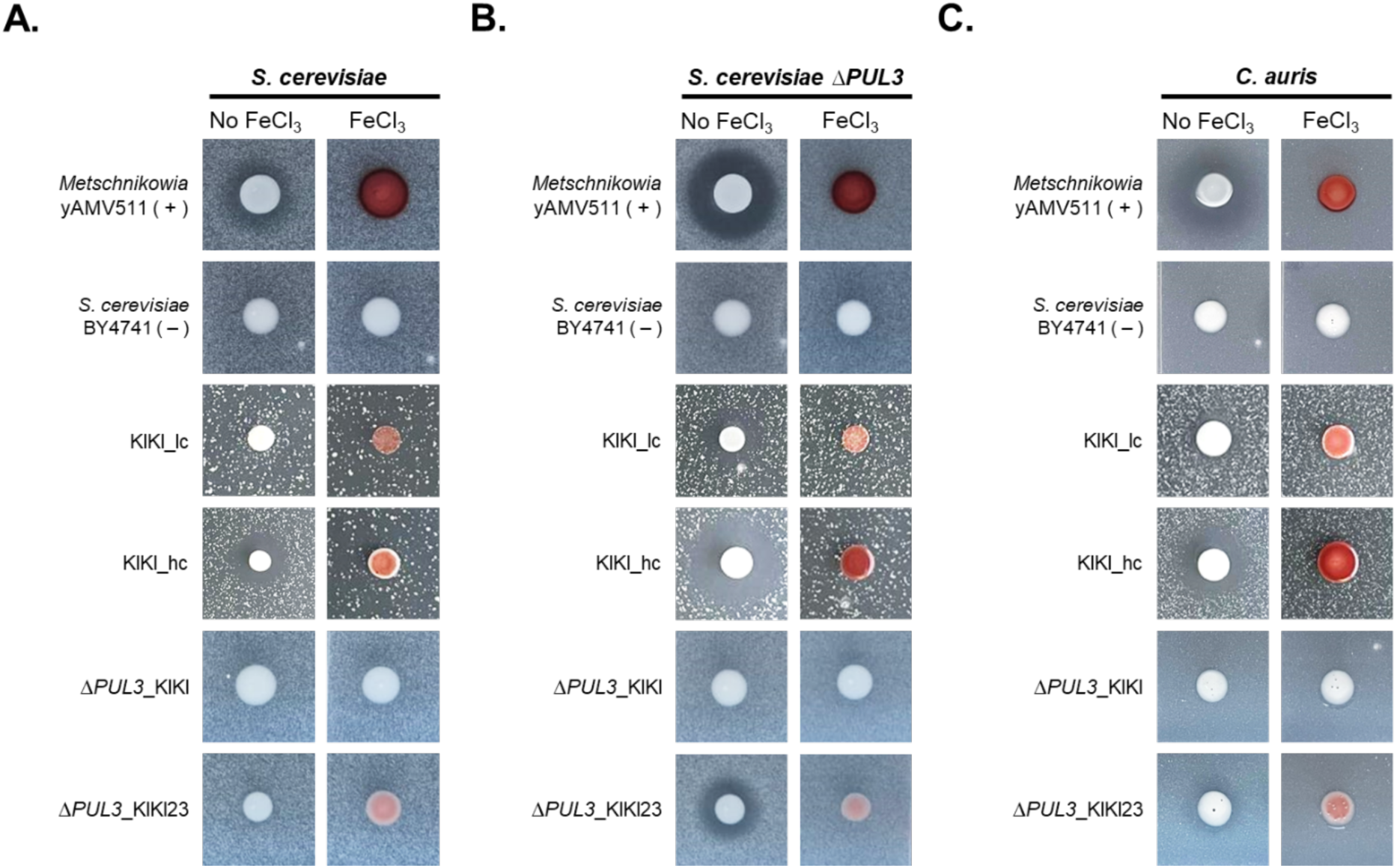
Growth inhibition assays with dual readout to explore the transport function of Pul3: cellular pigmentation for pulcherriminic import in media with 20 µg/mL FeCl_3_ and growth inhibition halo for pulcherrimin import. Pictures are representative of results from assays performed in triplicate. Plates were incubated at 30°C for 48 hours. The positive control was wild isolate yAMV511 (+) and the negative control was *S. cerevisiae* co– transformed with pRS413 and pRS416 empty plasmids (–). Cells of **(A)** *S. cerevisiae,* **(B)** *S. cerevisiae* Δ*PUL3* and **(C)** *C. auris* were embedded in agar and used as target strains.

In summary, we show a fully functional heterologous pulcherriminic acid production system, that also replicates the iron–dependent growth inhibition. However, high expression is required to achieve growth inhibition. Our findings complement previous results where *PUL* deletion studies in different *Metschnikowia* strains had shown loss of fungal growth inhibition against the filamentous phyto–pathogens *Geotrichum citri*–*aurantii* [27] and *Botrytis cinerea* [29].

### Expression of Pul3 reduces the sensitivity of non**–**producers/utilizers (cheaters) and is essential for producer strains to secrete pulcherriminic acid and inhibit fungal growth

In this study, we showed that *S. cerevisiae* – even though expressing a functional *PUL3* – was still sensitive to pulcherriminic acid–based iron deprivation. This provided an opportunity to explore two key aspects of Pul3 function as a transporter. First, we explored the previously hypothesized role of Pul3 as an importer [10] in the context of growth competition with non– producer organisms both utilizers (cheaters) and non–utilizers. Second, we investigated whether the transport function of Pul3 extends beyond import to the export of pulcherriminic acid. We studied those two aspects of Pul3 transport function making use of our functional heterologous *PUL* system in *S. cerevisiae* and assays to separately report on import (maroon pigmentation in iron–supplemented media) as well as export of pulcherriminic acid (presence of a halo when tested in media with trace amounts of iron).

Pul3 was first described by Krause and colleagues [10] as a putative transporter necessary for the uptake of pulcherrimin and reutilization of iron inside of cells expressing this protein. Given that *S. cerevisiae* contains a functional endogenous *PUL3* gene that allows it to uptake pulcherrimin and that it showed sensitivity against our isolates [15], we used *S. cerevisiae ΔPUL3* from the yeast deletion collection [46] as a target strain and compared its susceptibility to that of *S. cerevisiae*. Our reasoning was that the lack of *PUL3* would prohibit pulcherrimin uptake and render cells more sensitive to the iron monopolization by a producer strain. Thus, we performed a Halo assay with strain KlKl_lc, which did not produce a halo above, and KlKl_hc, which previously showed an inhibitory halo (**Figure 3A**). We found *S. cerevisiae ΔPUL3* to be more sensitive to pulcherriminic acid: the strain KlKl_lc did not display a growth inhibition halo against *S. cerevisiae* but did against *S. cerevisiae ΔPUL3* (**Figure 3A and 3B**); while strain KlKl_hc showed expected growth inhibition against both target strains but a bigger inhibitory halo against *S. cerevisiae ΔPUL3.* The distinct phenotypes confirmed that *PUL3* in *S. cerevisiae* is functionally involved in the uptake of extracellular pulcherrimin (produced by the producer cells) and able to reutilize the bound iron. Moreover, these results indicate that Pul3 is not sufficiently active (or expressed) to render the cells completely immune against the iron–depletion inhibitory mechanism. The use of a low– and high–production strains (KlKl_lc and KlKl_hc, respectively) provided the opportunity to study different levels of sensitivity and discover that low producers of pulcherriminic acid can inhibit non–producers but high producers are required to outcompete cheater organisms.

Next we studied if, besides pulcherrimin uptake, Pul3 was also necessary for the export of pulcherriminic acid, similarly to its bacterial equivalent Yvma in *B. licheniformes* [47]. We tested this theory with a deletion and complementation approach by transforming *S. cerevisiae ΔPUL3* with the low–copy plasmids encoding the *K. lactis PUL1* and *PUL2* genes – generating strain *ΔPUL3*_KlKl – and screened for growth inhibition using our Halo assay, as the production of a halo and maroon pigmentation would indicate pulcherrimin export. Neither growth inhibition nor maroon pigmentation in iron–supplemented media were observed against *S. cerevisiae* (**Figure 3A**). We used *S. cerevisiae ΔPUL3* for higher sensitivity but the same phenotype was observed (**Figure 3B**), meaning that *PUL3* encodes for a protein that is needed for the secretion of pulcherriminic acid and hence growth inhibition. We then confirmed this result by complementing the *PUL3* deletion by a plasmid encoded copy of *PUL3* from *K. lactis* (strain *ΔPUL3*_KlKl23), which recovered both pigmentation and the inhibitory activity against *S. cerevisiae ΔPUL3* (**Figures 3A and 3B**).

### Pulcherriminic acid inhibits the growth of the pathogenic fungus *C. auris*

Given that even a *PUL3*–encoding cheater such as *S. cerevisiae* was inhibited by our high–copy heterologous producer (although less sensitive than the *ΔPUL3* non–cheater strain), we wondered if our heterologous system would also inhibit a “potential cheater” like the pathogenic *C. auris,* which encodes for the complete four–gene *PUL* cluster **(Figure 1B).** In our previous work, we had shown that our iron–dependent growth inhibitory wild yeast isolates (including yAMV511) were active against this fungal pathogen [15]. Our heterologous *PUL* system would allow us to explore whether this inhibitory phenotype was solely due to pulcherriminic acid production.

Using the same Halo assay, we tested strains KlKl_lc and KlKl_hc against embedded *C. auris* cells. Interestingly, even the low–copy strain KlKl_lc showed certain growth inhibition against *C. auris*. The high–copy strain KlKl_hc showed a visually larger inhibitory halo (**Figure 3C**), as previously seen against *S. cerevisiae*. This meant that our high–production strains were able to inhibit the growth of this organism and that pulcherriminic acid production alone was a sufficient antifungal trait that could be transferred to other yeast.

In agreement with previous reports, we did not observe any maroon pigmentation of *C. auris* when grown in media supplemented with FeCl_3_, indicating that this species – despite encoding for the biosynthesis genes *PUL1* and *PUL2* – showed no visible pulcherriminic acid production [10]. It is possible that *C. auris PUL1* and/or *PUL2* are either not functional or not expressed under laboratory conditions, making it also questionable if that is the case for *PUL3* as well. It could be that *C. auris PUL* cluster is cryptic, as suggested by Krause and colleagues [10], and hence does not act as a cheater against our heterologous system in our assay. Alternatively, it could be that only *PUL1, PUL2* or both are no longer functional, while *PUL3/4* remain active, representing a new genotype for phenotypical cheaters, as well as an intermediate evolutionary step between producers and cheaters. Further studies on *C. auris* and other organisms with a complete *PUL* gene cluster but no pulcherriminic production, such as *Kluyveromyces dobzhanskii* and *Kluyveromyces wickerhamii* [10], would help clarifying on the functionality of *PUL3* in these type of potential cheaters.

However, the fact that the low–copy strain KlKl_lc was able to inhibit *C. auris* but not *S. cerevisiae* hints that *PUL3* might indeed not be active in this organism at least under the conditions used in our assay. Further research would be needed to verify if *C. auris PUL3* is expressed under different laboratory conditions and indeed functional.

### The cheating paradigm: competition and antagonism between pulcherriminic acid producers, utilizers and non**–**producers

Krause and colleagues first reported the *PUL* cluster and presented a model featuring three different phenotypes that a given yeast isolate can exhibit, based on their capacity for production and reutilization of pulcherriminic acid, which is dependent on the presence and expression of the *PUL* gene cluster or parts thereof in their genome [10]. We now extend this model by incorporating the inhibitory profile and the susceptibility profile of these three yeast phenotypes, based on our results on the function of Pul3. This can guide a better understanding of pulcherriminic acid–based competition, especially in the light of biocontrol or medical applications.

We observed three phenotypes: First, there are “producers” that encode for a complete functional *PUL* gene cluster (*PUL1–4*) and hence can synthesize pulcherriminic acid (*PUL1/2*), secrete it (*PUL3*), uptake and reutilize it (*PUL3*) and inhibit other yeast. Second, there are “non– producers/non–utilizers” that do not encode any *PUL* genes and therefore do not produce or reutilize pulcherriminic acid, hence are susceptible to pulcherriminic acid–based iron depletion. Third, there are “cheaters” (or “non–producers/utilizers”) that do not produce pulcherriminic acid but are able to uptake pulcherrimin and utilize the iron in their metabolism (*PUL3*).

In **Figure 4**, we show how the growth of non–producers/non–utilizers and non– producers/utilizers (cheaters) is affected by the presence of pulcherriminic acid producers. In the presence of an organism that produces low concentrations of pulcherriminic acid (low– producer; *e.g.* strain KlKl_lc), the growth of a non–producer is inhibited by iron depletion, as the element gets locked away and unavailable for its use (**Figure 4A**). If a producer produces higher concentrations of pulcherriminic acid (high–producer; *e.g.* strain KlKl_hc), the non– producer is still inhibited and a bigger inhibitory halo is seen in tests like our Halo assay, as the iron chelator diffuses and the minimal inhibitory concentration is reached further from the spot of producer cells (**Figure 4B**).

**Figure 4.**
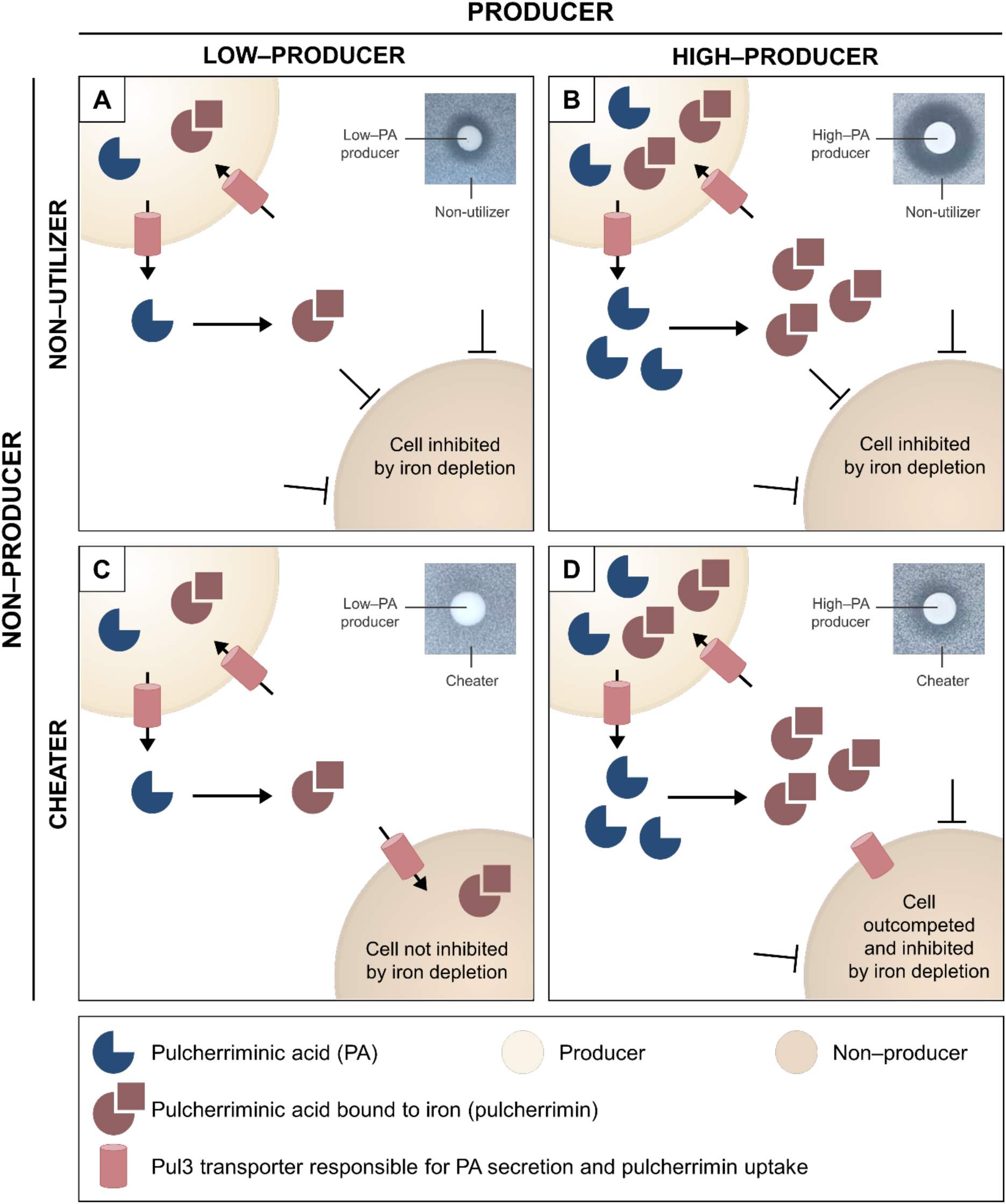
Model on the antagonistic interaction between organisms producing pulcherriminic acid (producers) and non–producers. Non–producers are divided in non–utilizers, unable to uptake iron–bound pulcherrimin, and utilizers/cheaters that produce Pul3 transporters for pulcherrimin uptake and reutilization). Each panel shows on the top right corner the readout of a Halo assay when testing both yeast phenotypes for growth inhibition. **4A.** Non–utilizers are inhibited via iron depletion. **4B.** Non–utilizers are strongly inhibited via iron depletion due to higher concentration of pulcherriminic acid bound to iron in the extracellular. **4C.** Cheaters produce Pul3 transporter, are able to uptake iron–bound pulcherrimin and are not inhibited against low–producers. **4D.** Cheaters produce Pul3 transporter, are able to uptake iron–bound pulcherrimin but get outcompeted by high–producers of pulcherriminic acid, eventually showing inhibition via iron depletion. The competitive advantage is hypothesized to rely on physical distance and difference in initial growth density but further research is needed.

The scenario looks different for the interaction between producers and cheaters: when a low– producer interacts with a cheater, no inhibition is seen with our Halo assay, meaning that the cheater can uptake pulcherrimin present in the extracellular and reutilize the iron bound (**Figure 4C**). Interestingly, and counterintuitively, we found that high–producers can still outcompete and inhibit the growth of cheater organisms (**Figure 4D**) and therefore potentially still be used as antifungals against *PUL3*–containing yeasts (cheaters).

Specifically, we tested the inhibitory activity of a high producer (KlKl_hc) against the cheater *S. cerevisiae* (**Figure 3**). In this setting, the producer and cheater are both *S. cerevisiae*, encoding the same *PUL3* gene. The constitutive expression of *PUL1/2* from *K. lactis* in the producer is the sole difference between them. Intuitively, a higher concentration of pulcherriminic acid in the environment should simply result in the cheater taking up a higher concentration of pulcherrimin, thus not being affected by the producer. However, our assay shows how the producer strain inhibited the growth of the cheating organism. We theorize that the inhibition is due to physical proximity between the producer cells and the high concentrations of iron that is sequestered in the form of pulcherrimin as compared to the distance to cheater cells. The producer organism produces pulcherriminic acid continuously, so that pulcherrimin is available in its immediate surroundings, while the cheater is dependent on pulcherriminic acid synthesis, secretion and diffusion from another organism. The proximity confers a competitive advantage that creates a local iron depletion zone that – although smaller when compared to the inhibition of non–producers – eventually does inhibit cheater cells. Besides, the setup of our halo assays leads to a growth advantage for producers, as the producer spot contains 5000–fold more cells than the embedded total amount of cheaters. Further research is needed on the ecological interactions of producers, non–producers and cheaters, and conditions on which the antagonism takes place, as well as the regulation and kinetics of the Pul3 transporter. A clear understanding on the nature of this competition will be essential for the use of pulcherriminic acid producers as biocontrol agents.

## Conclusions

In this study, we did not find any novel species producing pulcherriminic acid but we showed the abundance of *Metschnikowia* species among yeast isolates inhibiting the growth of other fungi by iron depletion. We also showed the level of sequence conservation in the *PUL* gene cluster, which allowed us to design primers for its detection in *Metschnikowia* isolates.

We heterologously expressed *PUL1 and 2* from *Metschnikowia* isolate yAMV511 and from *K. lactis* in *S. cerevisiae* and assessed pulcherriminic acid production using a pigmentation assay. Although *PUL1 and 2* are conserved across yeast, the expression of the yAMV511 genes only resulted in visible pulcherriminic acid production in a high copy vector, with *PUL1* being the production bottleneck. In contrast, genes from *K. lactis* showed cellular pigmentation, indicative of pulcherriminic acid production, even when using a low–copy expression system.

The heterologous expression of *PUL1/2* from *K. lactis* in *S. cerevisiae* enabled us to characterize the production, transport, and inhibitory activity of pulcherriminic acid in a “clean” heterologous background strain. We showed that the expression of those two genes, when expressed in a *PUL3–*encoding organism, were sufficient to transfer pulcherriminic acid production and growth inhibition of different yeast, including the major pathogen *C. auris* and *PUL3*–expressing cheaters like *S. cerevisiae*. Moreover, we showed that Pul3 is responsible for both the export of pulcherriminic acid to the extracellular and import of iron–bound pulcherrimin and hence essential for yeast to exert their inhibitory activity.

### Future perspectives on fundamental research

Our functional heterologous system can enable further characterization of the proteins involved in the biosynthesis of pulcherriminic acid. For example, we identified several residues in both Pul1 and Pul2 that are conserved across *Metschnikowia* isolates and *K. lactis* (**Supplementary Figures 1 and 2**). Those conserved residues might be important for enzymatic catalysis or structural stability of both proteins. For instance, within the conserved positions, Pul1 presents a glycine–rich region (positions 236–243 in *K. lactis;* essential for binding to nucleotides [48]); while Pul2 contains a **WxxxR** motif (positions 145–149 in *K. lactis*; heme–binding conserved sequence in P450 cytochromes [49]). The generation of scanning libraries for both enzymes, possibly using these conserved residues highlighted as a started point, would provide valuable information on which positions and residues are important for substrate binding or activity [50]–[52]. Furthermore, assays on cellular pigmentation in iron–supplemented media could serve as a straightforward screening method for loss of functionality.

Another approach to elucidate essential residues would be the study on locus *PUL1* from *M. pulcherrima* KIOM G15050 and *PUL2* from *M. pulcherrima* AP4, which contained stop codons in their sequence and showed two ORFs for two small proteins instead of one like the rest of *Metschnikowia* strains (**Figure 1A**). When aligned to the rest of Pul proteins, those showed sequence similarity to distinct parts of other Pul proteins with no overlapping regions (**Supplementary Figures 1 and 2**). It seems that those yeasts produce two different proteins rather than multiple copies of the same protein present in the same cluster as previously suggested for *M. pulcherrima* KIOM 15050 [44]. Since both strains have been reported to effectively produce pulcherriminic acid [21], [53], studies with constructs expressing only one of those two smaller proteins would help elucidating whether there were false stop codons introduced when performing the whole–genome shotgun sequencing, only one of the two proteins is sufficient for its function on the biosynthesis of pulcherriminic acid, or whether both proteins are expressed as two subunits that interact *in vivo* and perform their enzymatic function similarly to the rest of pulcherriminic acid producers. In the case that only one protein was sufficient, the residues present in the other small protein would be automatically described as dispensable.

### Application of our system for the development of novel antifungals

Pulcherriminic acid–producing yeasts previously showed inhibitory activity against human pathogens including *N. glabratus, C. auris* and other *Candida* species [15], [19].

Some bacteria produce sideromycins, naturally occurring conjugates of siderophores and antibiotics, such as albomycin, ferrimyicin and salmycin [6], [54]. This innate targeting strategy commonly known as ‘Trojan horse’ is naturally used by producers to scavenge for iron and inhibit competitors [55]–[57]; and it recently inspired the research community to synthesize similar compounds both in bacteria [58]–[62] and fungi [63], [64].

The conjugation of small antifungal compounds to pulcherriminic acid would be of special interest to develop novel antifungals against human pathogens, as the biosynthesis of this iron chelator, in comparison with siderophores commonly used for such conjugates, is not restricted to iron–depleted conditions. Moreover, the conjugation of pulcherriminic acid to antifungals would provide drugs with certain specificity towards target organisms with functional Pul3 receptors.

### Application of our system for biocontrol

The potential of yeasts producing pulcherriminic acid as biocontrol agents against phyto– pathogens has been discussed [17], [18], [65] and already led to the development of several products for bioprotection of harvest fruits and prevention of the spoilage of fermented drinks. These organisms previously showed inhibitory activity against plant pathogens like *Botrytis, Geotrichum, Penicillium* and *Alternaria* [18], [27] but also against food spoilage yeasts like *Brettanomyces and Zygosaccharomyces* [15], [17], [19].

Some examples of products in the market based on pulcherriminic acid–producing yeasts are Excellence® B–nature® from Lamothe–Abiet©, Gaïa™ from Lallemand©, Noli® from Koppert© and Zymaflore® Khio from Laffort©. However, better understanding on the molecular basis of the expression, regulation and production is necessary for their adequate use. In fact, some yeast–based products have been withdrawn from the market before due to ineffective or inconsistent results under commercial conditions; for example, Candifruit® [66], Aspire® [67] and Yieldplus® [68].

For biocontrol applications, we observed that natural *Metschnikowia* species produce higher concentrations than our heterologous system, based on pigmentation and growth inhibition assays. Studies on growth optimization, like in *Metschnikowia pulcherrima* CBS 10809 (previously *M. andauensis*) [69] could increase pulcherriminic acid production; however, strain optimization requires fundamental knowledge on the expression, regulation and mode of action of pulcherriminic acid. In this context, our heterologous system in a clean background confirmed that the sole production of this iron chelator is sufficient to inhibit the growth of other organisms. Likewise, our system – built in such a well–characterized organism as *S. cerevisiae* – will allow further studies on remaining unknown aspects separately; for example, with further research on the functionality of *PUL4* as regulator of the *PUL* gene cluster.

## Acknowledgements

The authors would like to thank the Center for Information Technology of the University of Groningen for providing access to the Hábrók high performance computer cluster.

## Contributions

AMV and SB designed the research. AMV, JJvW and AJF performed the experiments. ARO and CM designed and performed the LC–MS experiments and analysed the data. AMV drafted the manuscript. SB revised the original manuscript. All authors have revised and approved the final version of the manuscript.

