## Supplementary information for "Pulcherriminic acid biosynthesis and Transport: Insights from a heterologous system in *Saccharomyces cerevisiae*"

**Supplementary Table 1**. Strains used in this work.

| Name | Characteristics and use | Source |
|---|---|---|
| *S. cerevisiae* BY4741 (–) | BY4741 pRS413 + pRS416; Negative control for pulcherriminic production and target strain for fungal growth inhibition | This work |
| MpMp_lc | BY4741 pPUL1Mp_lc + pPUL2Mp_lc; pulcherriminic acid production in low copy plasmids with *PUL1* and *PUL2* genes from yAMV511 | This work |
| MpMp_hc | BY4741 pPUL1Mp_hc + pPUL2Mp_hc; pulcherriminic acid production in high copy plasmids with *PUL1* and *PUL2* genes from yAMV511 | This work |
| KlKl_lc | BY4741 pPUL1Kl_lc + pPUL2Kl_lc; pulcherriminic acid production in low copy plasmids with genes from *K. lactis* | This work |
| KlKl_hc | BY4741 pPUL1Kl_hc + pPUL2Kl_hc; pulcherriminic acid production in high copy plasmids with genes from *K. lactis* | This work |
| MpKl | BY4741 pPUL1Mp_lc + pPUL2Kl_lc; pulcherriminic acid production with low copy plasmids with *PUL1* from yAMV511 and *PUL2* from *K. lactis* | This work |
| KlMp | BY4741 pPUL1Kl_lc + pPUL2Mp_lc; pulcherriminic acid production with low copy plasmids with PUL1 from *K. lactis* and PUL2 from yAMV511 | This work |
| *S. cerevisiae* Δ*PUL3* | BY4741 Δ*PUL3* empty plasmid 1 + empty plasmid 2; Target strain for fungal growth inhibition | This work |
| Δ_KlKl | BY4741 Δ*PUL3* pPUL1Kl + pPUL2Kl; pulcherriminic acid production with low copy plasmids with genes from *K. lactis* | This work |
| Δ_KlKl23 | BY4741 Δ*PUL3* pPUL1Kl + pPUL2Kl_PUL3Kl; pulcherriminic acid production with low copy plasmids with genes from *K. lactis* | This work |
| *C. auris* ATCC-MYA-5001 | Target strain for fungal growth inhibition | American Type Culture Collection |
| yAMV240 | Wild isolate - species identification; *PUL1* and *PUL2* sequencing | [1] |
| yAMV312 | Wild isolate - species identification; *PUL1* and *PUL2* sequencing | [1] |
| yAMV460 | Wild isolate - species identification; *PUL1* and *PUL2* sequencing | [1] |
| yAMV511 | Wild isolate - species identification; *PUL1* and *PUL2* sequencing; pulcherriminic acid production via LC-MS | [1] |
| yAMV636 | Wild isolate - species identification; *PUL1* and *PUL2* sequencing | [1] |
| yAMV642 | Wild isolate - species identification; *PUL1* and *PUL2* sequencing | [1] |
| yAMV32 | Wild isolate - species identification | [1] |
| yAMV41 | Wild isolate - species identification | [1] |
| yAMV99 | Wild isolate - species identification | [1] |
| yAMV160 | Wild isolate - species identification | [1] |

| | | |
|---|---|---|
| yAMV174 | Wild isolate - species identification | [1] |
| yAMV186 | Wild isolate - species identification | [1] |
| yAMV204 | Wild isolate - species identification | [1] |
| yAMV215 | Wild isolate - species identification | [1] |
| yAMV217 | Wild isolate - species identification | [1] |
| yAMV233 | Wild isolate - species identification | [1] |
| yAMV286 | Wild isolate - species identification | [1] |
| yAMV322 | Wild isolate - species identification | [1] |
| yAMV346 | Wild isolate - species identification | [1] |
| yAMV360 | Wild isolate - species identification | [1] |
| yAMV380 | Wild isolate - species identification | [1] |
| yAMV420 | Wild isolate - species identification | [1] |
| yAMV564 | Wild isolate - species identification | [1] |
| yAMV610 | Wild isolate - species identification | [1] |
| yAMV623 | Wild isolate - species identification | [1] |
| yAMV660 | Wild isolate - species identification | [1] |
| yAMV669 | Wild isolate - species identification | [1] |
| yAMV692 | Wild isolate - species identification | [1] |
| yAMV702 | Wild isolate - species identification | [1] |
| yAMV721 | Wild isolate - species identification | [1] |

**Supplementary Table 2.** Oligonucleotides used in this study for species identification, sequencing of *PUL* cluster in wild yeast isolates and molecular cloning of *PUL* genes in a *S. cerevisiae* heterologous system.

| **Name** | **Description** | **Source** | **Sequence (5' → 3')** |
|---|---|---|---|
| Species identification | | | |
| ITS3 (fw) | ITS2 region from wild yeast isolates | [2] | GCATCGATGAAGAACGCAGC |
| ITS4 (rv) | ITS2 region from wild yeast isolates | [2] | TCCTCCGCTTATTGATATGC |
| NL1 (fw) | D1/D2 domain from wild yeast isolates | [3] | GCATATCAATAAGCGGAGGAAAAG |
| NL4 (rv) | D1/D2 domain from wild yeast isolates | [3] | GGTCCGTGTTTCAAGACGG |
| *PUL* cluster identification and sequencing | | | |
| PUL1fw | *PUL1* from wild yeast isolates | This work | AGAATACAGGTGGGCTCA |
| PUL1midfw | *PUL1* from wild yeast isolates | This work | ATGAAATGATATGCACCCACA |
| PUL1rv | *PUL* cluster from wild yeast isolates | This work | TTCTAGATAAAGAGAACACCCTGTT |
| PUL2midrv | *PUL2* from wild yeast isolates | This work | TTTATCGTTGGCCATGGTG |
| PUL2rv | *PUL2* from wild yeast isolates | This work | AATTGGTTTCCTAATCGGGA |
| PUL3fw | *PUL* cluster from wild yeast isolates | This work | TTACAGTTACATCACAGCCATAC |
| *Molecular cloning* | | | |
| SB618 | *PUL1* fw from *K. lactis NRRL Y-1140* | This work | GCATCGTCTCATCGGTCTCATATGTACCAACTGCTTTTCC |
| SB619 | *PUL1* rv from *K. lactis* NRRL Y-1140 | This work | ATGCCGTCTCAGGTCTCAGGATTCAGATTACGAGAGCACCA |
| SB620 | *PUL2* fw from *K. lactis NRRL Y-1140* | This work | GCATCGTCTCATCGGTCTCATATGTTAGCTGATATATTAATCCCA |
| SB621 | *PUL2* rv from *K. lactis* NRRL Y-1140 | This work | ATGCCGTCTCAGGTCTCAGGATTCACAATGCAGTTAGTT |
| SB622 | *PUL3* fw from *K. lactis NRRL Y-1140* | This work | GCATCGTCTCATCGGTCTCATATGAAGCTAACAGATTCACAAA |
| SB623 | *PUL3* rv from *K. lactis* NRRL Y-1140 | This work | ATGCCGTCTCAGGTCTCAGGATCACATTTTGTTCCTCCTAAG |
| SB626 | *PUL3* fw (BsaI outcloning) | This work | GCATCGTCTCAGAGCGGTATCTTTAAACACGTTACTTG |
| SB627 | *PUL3* rv (BsaI outcloning) | This work | GCATCGTCTCAGCTCCTGCACAGGAATATA |
| SB628 | *PUL2* fw (BsaI outcloning) | This work | GCATCGTCTCAGAGATCAGTGTGGTGACCTG |
| SB629 | *PUL2* rv (BsaI outcloning) | This work | GCATCGTCTCATCTCTGTTTTTGATAGACAAGCTTTTGCCG |

**Supplementary Table 3.** DNA sequence of *PUL1/2* from yAMV511 ordered as gBlocks. The shaded part of the sequence indicates the overhangs necessary to clone the oligonucleotides into plasmids following the MoClo system into Type 1 plasmids [4].

| | |
|---|---|
| PUL1Mp | GCATCGTCTCATCGGTCTCATATGGTGACTGAATCAGGCGAGCTGTTCATAAGCCAGGCCT<br>TCATAGACATGTGGATAGATAGCCTAGTTGGTTATCTACCCGGAGAAACGTGCAGACACGT<br>TTCTCAATTTGTCCAGAATGAGTACGTGGGTACGCTGGGTCAATTATACGTCACCATAAGG<br>GACTTGAGGCAGGCCATCAGCGCATTCGTGGACAGTGTTAATGGCGAAGAAAATAAAAATT<br>TTTTGGCCTTTGATGCACATGTATCTGACTGTGGTTGCCATTTGCGTGCGTCCATGAGTAT<br>GGACTTGATTCAAAGATACAGGGGCAATAGGGAAGAGTTACTGAGTTTTCTAGGCTTGGTC<br>GAAGCTTGCGATAGGGCTCTGGTCAGTACAAGTGCCTTAATGAAGGATATATGCACTGAGG<br>CGAAAAGCTTGAAAGAATTACAACTACCTAAAAGTACCAAAGATCCTTTACTGTTTTTGAA<br>CGCAATCGGCTGGAAATTTGAATCTAACAACTTGAGTGAAATAAAATATATCTTCTACTGC<br>TACGTGTTATCACAATTCAAGACGTATAGCTTCAGGAACAAACAAGATTCAGTCCACATTG<br>ATACAGACAAAGAATTTAAACAGAAGAATGAAATGATTTGTACTCACACCTGTCAGGGCAA<br>GGGAAAGCTGGGAAATGGGTGTAGATATCTGAAGCACGCGAGGATTGGGAAGGCTGCGCTG<br>AAACAATGGACACTTTGTTATCAGGAGAGGCTGTCCAAAATGTCCGTGGACTATCTTGCTA<br>AGTCAGACTCAGAGTTGAAAGAGCTGGTAGAGACAAGCCGTAAAGAATCCCATAAGTCTGT<br>CGCAGCAGTACCGTCTTATGTCCAATTTAAAATATCCGAGCGTCTGTGGGCGAGTAATCAA<br>TTCCCATTTCTTCTTAGTATGAGAGTTTTTGTCGATGAGGGGCACGACGATATATACGCCC<br>GTGCCTTCGTCGGTCGTGATCTTAAATGGAACATCCAATTCGTAACATCAGACGTCTTAGA<br>GGATACGCCACACATTATCGTGGCAGGGCATTGTCGTGTACCCCACGGCTACAATGACATA<br>AACAAGTTAGACCTAGCTAACTTGTCACTTGATGCTCATCAAAATATGAGGAGTTTTTGGT<br>ATTCATTTATGTCACAACACAAACAGTACCCCTTTGACACCGCCCTAGGCTGTGATGATGA<br>CCTTCAAAATGTTTTACCTGCCCATGAATTTAAGGATTACATGAAGTTTAAGTATGCCGGC<br>ATAAGGGCATTTCGTGATATGGAATTCACACCAAAGCACATTTTTGTTGAATATCCAAGTA<br>TCGTGTTCAGTAAGCAGAGAATGTTGGCTGGGAAACAGGGCGTCCTATTCATTTAGATCCT<br>GAGACCTGAGACGGCAT |
| PUL2Mp | GCATCGTCTCATCGGTCTCATATGCTAACCATACTATCCCTTCCGGTCTTTTGGGTACTTG<br>TAGTTAGTTTGTGTCTAGTCAGTTCCAAAAGTAAACTTAGGGCCTTTCAAAAGCAACCACC<br>ATTTAAGAAGATACCCAGATCCCATCGTAACGTAGAGGGAAAGAAAATTACACAGGGTCCG<br>GAGATCTCTGAGAAAAACAAAAATCAGTACGGCAGCATATATTCACATCGTGATGGGTTTA<br>GATATGAGGTCGTTCTAACAACGCCTAGCCAATTAAAACAATACTACAGCTCTCATACGAA<br>AGATCACAAGAAACTGGATAGCTTTGGAGCAGGGCAGTATCTTGTCGCGCTTCTGGGGGAA<br>TGTCTTGGGTTCCAAAACGGGGAGAGTTGGACCAGAATGAGAAAGAGTTTTAACTTGTTTT<br>TCACCCATACCTTAGCCGCTAAGACACTTCCTGCAATGATAGCGTTCATCGACGGATGGAT<br>TTCTGAGCATGACAGCCAGAATGAATTCAGTGTAGACGCCTTCGACTTCGTTGCAACCGTA<br>CCATTTACTTGCATTGCCAAGTATCTGTACGGAAACGAGCAGTGCAGCGGGCGTGTCCTGA<br>CTGAGCTTAAGAACCTTGTACCTTTGCACAGTGAACTTATGACTCACGCCTTTACCACTTT<br>CTGGGGTCGTTTTCGTATCTACCAATACTTCCCTTTTCAGAGGATGAAGGATTTGAAGTAT<br>TTTCAAGATTCTTTCAAGTCACTTAGCCTAGCAATGGTCGAGTCAGCCCGTGACGCCGAGA<br>ACCCAACTGTTGCTTCCGAATTGTACAAGTTAGTCGAAAGCCGTGATTTGACGCTTGATAA<br>TTGGATACAATCCCTAGACGAGATTCTGTTCGCCAATATCGATGTTACTGCTACGATAATG<br>TCTTGGTCATTGGTTGAGATGGGTCGTAACAAACACGAGCAAGCCCGTCTGAGGCTTGAGG<br>TACTGGAGAACCTTCAGTCAGTAGACGAATATTGCAAAAGAACAGACACTGTTCTACATAG<br>AGTCTTACTAGAGATCCTTAGGCTTCATCCACTATTGTGGTATGGTTTCCCCGAACAAAGT<br>AGCAGTGCAATGGTTATCGATGGTCACAAGATCGAGGCTAACACTCCCATTGTGGTTGATC<br>AGTACCAATTAAATTACAAGAGCCCATTATGGAACCCAGCGGATAAGAGCACGGACTATGG<br>AGCAACTTTCGATTCCAACCGTTTTCTAGGACTTAATAATCGTGATATATTGATGTCCTCT<br>GTAACGTTTGGGAGCGGGCCTCGTAGGTGCTTAGGGAAAAACTTCGCCGAAGTTCTGATCA<br>AGACGGAAGTGGCGAAGGTGTTGAGCACATTCGAGGTCGCGCTTGAAGGGGAGTTAAAGGT<br>AGCTGCCGACACTTTCGTTGTAAGACCCGATGCGCAAATAAAACTTACAAGGTTGATATAG<br>ATCCTGAGACCTGAGACGGCAT |

**Supplementary Table 4.** Plasmids used in this work.

| **Name** | **Characteristics and use** | **Source** | **Link** |
|---|---|---|---|
| *Yeast Toolkit parts* | | | |
| pYTK001 | Entry vector | [4] | https://www.addgene.org/65108/ |
| pYTK002 | conLS (Type 1) | [4] | https://www.addgene.org/65109/ |
| pYTK009 | pTDH3 (Type 2) | [4] | https://www.addgene.org/65116/ |
| pYTK010 | pCCW12 (Type 2) | [4] | https://www.addgene.org/65117/ |
| pYTK014 | pTEF2 (Type 2) | [4] | https://www.addgene.org/65121/ |
| pYTK047 | GFPdropout (Type 234r) | [4] | https://www.addgene.org/65154/ |
| pYTK051 | tENO1 (Type 4) | [4] | https://www.addgene.org/65158/ |
| pYTK052 | tSSA1 (Type 4) | [4] | https://www.addgene.org/65159/ |
| pYTK053 | tADH1 (Type 4) | [4] | https://www.addgene.org/65160/ |
| pYTK072 | conRE (Type 5) | [4] | https://www.addgene.org/65179/ |
| pYTK074 | URA3 (Type 6) | [4] | https://www.addgene.org/65181/ |
| pYTK076 | HIS3 (Type 6) | [4] | https://www.addgene.org/65183/ |
| pYTK081 | CEN6/ARS4 (Type 7) | [4] | https://www.addgene.org/65188/ |
| pYTK082 | 2 micron (Type 7) | [4] | https://www.addgene.org/65189/ |
| pYTK083 | AmpR-ColE1 (Type 8) | [4] | https://www.addgene.org/65190/ |
| pYTK084 | KanR-ColE1 (Type 8) | [4] | https://www.addgene.org/65191/ |
| pYTK095 | AmpR-ColE1 (Type 678) | [4] | https://www.addgene.org/65202/ |
| *Plasmids constructed* | | | |
| EP1 | pRS413: Negative control | This work | https://benchling.com/s/seq-Plf3Ui11zvNvyrxJ6DQI?m=slm-YA7sS7zLQobwxtyupcys |
| EP2 | pRS416; Negative control | This work | https://benchling.com/s/seq-0pguFzdsGasAf9ZlqMNI?m=slm-t7AUiC0YXFMxlocq58Wg |
| pPUL1Mp_lc | *PUL1* from *Metschnikowia* sp yAMV511 in low–copy plasmid | This work | https://benchling.com/s/seq-K5mBajixUn3DECn9A49d?m=slm-zIEXfcUJtRJ8CUZod67b |
| pPUL1Mp_hc | *PUL1* from *Metschnikowia* sp yAMV511 in high–copy plasmid | This work | https://benchling.com/s/seq-9R5BzrKMgaEQSRCXty6n?m=slm-kKUjea3azkQxgJXaX999 |
| pPUL2Mp_lc | *PUL2* from *Metschnikowia* sp yAMV511 in low–copy plasmid | This work | https://benchling.com/s/seq-ugapG7UYTtP2vWY7zXKi?m=slm-nn702CTEFPOD526sznRy |
| pPUL2Mp_hc | *PUL2* from *Metschnikowia* sp yAMV511 in high–copy plasmid 1 | This work | https://benchling.com/s/seq-robtQMtdM2HmgyfiHxxq?m=slm-ZEeS1R5ThNWoNpCt2nDS |
| pPUL1Kl_lc | *PUL1* from *K. lactis* in low–copy plasmid | This work | https://benchling.com/s/seq-YMPq7KStg046NbcNMw7s?m=slm-41YHDDBQMoIAyzHrxUpL |

| | | | |
|---|---|---|---|
| pPUL1Kl_hc | *PUL1* from *K. lactis* in high–copy plasmid | This work | https://benchling.com/s/seq-Z0RXnM2FkzIDvUAUudoF?m=slm-dcr3tYWtOgBo08qIQ4Og |
| pPUL2Kl_lc | *PUL2* from *K. lactis* in low–copy plasmid | This work | https://benchling.com/s/seq-naWL6n0lxD4FCT24kOo3?m=slm-VLNdBIEJqf04W3lyNJWO |
| pPUL2Kl_hc | *PUL2* from *K. lactis* in high–copy plasmid | This work | https://benchling.com/s/seq-l5EEp7OnGp0kIp29KoS5?m=slm-iacKt48pHGCWRXCM7Mvv |
| pPUL2Kl_PUL3Kl | *PUL2/3* from *K. lactis* in low–copy plasmid | This work | https://benchling.com/s/seq-xSrSiOCXK64DQ1wn3ZlW?m=slm-bCDzkUfN6OYWU1ZXIFTQ |

**Supplementary Table 5.** Species identification of yeast isolates producing iron chelators. The table shows BLAST hits with highest sequence similarity. Isolates were linked to a given species when 97% sequence similarity was reached and to a given genus when >95% was reached from the hits retrieved. All isolates belong to the *Metschnikowia* clade.

| yAMV # | OTU | Locus | Sequence (5' → 3') |
|---|---|---|---|
| 32 | *M. pulcherrima* | D1/D2 | TCAGTAACGGCGAGTGAAGCGGCAAAAGCTCAAATTTGAAATCCCCCGG<br>GAATTGTAATTTGAAGAGATTTGGGTCCGGCCGGCGGGGGTTAAGTCCA<br>CTGGAAAGTGGCGCCACAGAGGGTGACAGCCCCGTGAACCCCTTTAACG<br>CCCTCATCCCAGATCTCCAAGAGTCGAGTTGTTTGGGAATGCAGCTCTA<br>AGTGGGTGGTAAATTCCATCTAAAGCTAAATACCGGCGAGAGACCGATA<br>GCGAACAAGTACAGTGATGGAAAGATGAAAAGCACTTTGAAAAGAGAGT<br>GAAAAAGTACGTGAAATTGTTGAAAGGGAAGGGCTTGCAAGCAGACACT<br>TAACTGGGCCAGCATCGGGGCGGCGGGAAACAAAACCACCGGGGAATGT<br>ACCTTTCGAGGATTATAACCCCGGTCTCTATTTCCTTGTTGCCCCGAGG<br>CCTGCAATCTAAGGATGCTGGCGTAATGGTTGCAAGTCGCCCGTCT |
| | *M. pulcherrima* | ITS2 | CTTGCAGTAACGTGAATCATTGAATCTTCTGAACGCACATCTGCGTCTC<br>TCTCGGGGTATTCCCCAGGGCATGCGTGGGTGAGCGATATTTACTCTCA<br>AACCTCCGGTTTGGTCCTGCTTCGGCCTAATATCAACGGCGCTAGAATA<br>AGTTTTAGCCCCATTCTTTTTCCTCACCCTCGTAAGACTACCCGCTGAA<br>CTTAAGCATATCATAAAGCCGG |
| 240 | *M. pulcherrima* | D1/D2 | GAAGAGATTTGGGTCCGGCCGGCGGGGGTTAAGTCCACTGGAAAGTGGC<br>GCCACAGAGGGTGACAGCCCCGTGAACCCCCTCAACGCCCTCATCCCAG<br>ATCTCCAAGAGTCGAGTTGTTTGGGAATGCAGCTCTAAGTGGGTGGTAA<br>ATTCCATCTAAAGCTAAATACCGGCGAGAGACCGATAGCGAACAAGTAC<br>AGTGATGGAAAGATGAAAAGCACTTTGAAAAGAGAGTGAAAAAGTACGT<br>GAAATTGTTGAAAGGGAAGGGCTTGCAAGCAGACACTTAACTGGGCCAG<br>CATCGGGGCGGCGGGAAACAAAACCACCGGGGAATGTACCTCTCGAGGA<br>TTATAACCCCGGTCTCAATTTCCTCGCCGCCCCGAGGCCTGCAATCTAA<br>GGATGCTGGCGTAATGGTTGCAAGTCGCCC |
| | *M. pulcherrima* | ITS2 | CCCCGGGGTATTCCCCAGGGCATGCGTGGGTGAGCGATATTTACTCTCA<br>AACCTCTGGTTTGGTCCTGCTTCGGCCTAATATCAACGGCGCTAGAATA<br>AGTTTTAGCCCCATCCTTTTTCCTCACCCTCGTAAGACTACCCGCTGAA<br>CTTAAGCATATCATAAAAGCGGAGG |
| 286 | *M. pulcherrima* | D1/D2 | AAATCCCCCGGGAATTGTAATTTGAAGAGATTTGGGTCCGGCCGGCAGG<br>GGTTAAGTCCACTGGAAAGTGGCGCCACAGAGGGTGACAGCCCCGTGAA<br>CCCCCTCAACGCCCTCATCCCAGATCTCCAAGAGTCGAGTTGTTTGGGA<br>ATGCAGCTCTAAGTGGGTGGTAAATTCCATCTAAAGCTAAATACCGGCG<br>AGAGACCGATAGCGAACAAGTACAGTGATGGAAAGATGAAAAGCACTTT<br>GAAAAGAGAGTGAAAAAGTACGTGAAATTGTTGAAAGGGAAGGGCTTGC<br>AAGCAGACACTTAACTGGGCCAGCATCGGGGCGGCGGGAAACAAAACCA<br>CCGGGGAATGTACCTTTCGAGGATTATAACCCCGGTCTCAATTTCCATG<br>TTGCCCCGAGGCCTGCAATCTAAGGATGCTGGCGTAATGGTTGCAAGTC<br>GCCCGT |
| | *M. pulcherrima* | ITS2 | TTGAATCTTTGAACGCACATTGCGCCCCGGGGTATTCCCCAGGGCATGC<br>GTGGGTGAGCGATATTTACTCTCAAACCTCCGGTTTGGTCCTGCTTCGG<br>CCTAATATCAACGGCGCTAGAATAAGTTTTAGCCCCATCCTTTTTCCTC<br>ACCCTCGTAAGACTACCCGCTGAACTTAAGCATATCAATAAGCGGAGGA<br>A |
| 312 | *M. pulcherrima* | D1/D2 | TCCCCCGGGAATTGTAATTTGAAGAGATTTGGGTCCGGCCGGCGGGGGT<br>TAAGTCCACTGGAAAGTGGCGCCACAGAGGGTGACAGCCCCGTGAACCC<br>CTTCAACGCCCTCATCCCAGATCTCCAAGAGTCGAGTTGTTTGGGAATG<br>CAGCTCTAAGTGGGTGGTAAATTCCATCTAAAGCTAAATACCGGCGAGA<br>GACCGATAGCGAACAAGTACAGTGATGGAAAGATGAAAAGCACTTTGAA<br>AAGAGAGTGAAAAAGTACGTGAAATTGTTGAAAGGGAAGGGCTTGCAAG |

| | | | |
|---|---|---|---|
| | | | CAGACACTTAACTGGGCCAGCATCGGGGCGGCGGGGAGCAAAACCACCG<br>GGGAATGTACCTTTCGAGGATTATAACCCCGGCCCTTACTCCCATACTG<br>CCCCGAGGCCTGCATTCTAAGGATGCTGGCGTAATGGTTGCAAGTCGCC<br>CGTC |
| | *M. pulcherrima* | ITS2 | TTTGAATCTTTTGAACGCACTCTGCGCCCCGGGGTATTCCCAGGGCATG<br>CGTGGGTGAGCGATATTTACTCTCAAACCTCCGGTTTTGGTCCTGCTTC<br>GGCCTAATATCAACGGCGTCTAGAATAAGTTTTAGCCCCATTCTTCTTC<br>CTCACCCTCGTAAGACTACCCGCTGAACTTAAGCATA |
| 420 | *M. pulcherrima* | D1/D2 | GTGAGCGGCAAAAGCTCAAATTTGAAATCCCCCGGGAATTGTAATTTGA<br>AGAGATTTGGGTCCGGCCGGCAGGGGTTAAGTCCACTGGAAAGTGGCGC<br>CACAGAGGGTGACAGCCCCGTGAACCCCTTTAACGCCCTCATCCCAGAT<br>CTCCAAGAGTCGAGTTGTTTGGGAATGCAGCTCTAAGTGGGTGGTAAAT<br>TCCATCTAAAGCTAAATACCGGCGAGAGACCGATAGCGAACAAGTACAG<br>TGATGGAAAGATGAAAAGCACTTTGAAAAGAGAGTGAAAAAGTACGTGA<br>AATTGTTGAAAGGGAAGGGCTTGCAAGCAGACACTTAACTGGGCCAGCA<br>TCGGGGCGGCGGGAAACAAAACCACCGGGGAATGTACCTTTCGAGGATT<br>ATACCCCCGGTCTCTATTTCCTTGCTGCCCCGAGGCCTGCAATCTAAGG<br>ATGCTGGCGTAATGGTTGCAAGTCGCC |
| | *M. pulcherrima* | ITS2 | TTGAACTTTGCAGTAACGTTGAATCATTTGAAATCCTTTTGAACGCACA<br>TCTGCGCCCCGGGGTATTCCCCAGGGCATGCGTGGGTGAGCGATATTTA<br>CTCTCAAACCTCCGGTTTGGTCCTGCTTCGGCCTAATATCAACGGCGCT<br>AGAATAAGTTTTAGCCCCAGCCTTTTTCCTCACCCTCGTAAGACTACCC<br>GCTGAACTTAAGCATAT |
| 460 | *M. pulcherrima* | D1/D2 | AATTTGAAGAGATTTGGGTCCGGCCGGCGGGGGTTAAGTCCACTGGAAA<br>GTGGCGCCACAGAGGGTGACAGCCCCGTGAACCCCTTCAACGCCCTCAT<br>CCCAGATCTCCAAGAGTCGAGTTGTTTGGGAATGCAGCTCTAAGTGGGT<br>GGTAAATTCCATCTAAAGCTAAATACCGGCGAGAGACCGATAGCGAACA<br>AGTACAGTGATGGAAAGATGAAAAGCACTTTGAAAAGAGAGTGAAAAAG<br>TACGTGAAATTGTTGAAAGGGAAGGGCTTGCAAGCAGACACTTAACTGG<br>GCCAGCATCGGGGCGGCGGGAAACAAAACCACCGGGGAATGTACCTTTC<br>GAGGATTATAACCCCGGTCTCTATTTCCTCGCCACCCCGAGGCCTGCAA<br>TCTAAGGATGCTGGCGTAATGGTTGCAAGTCGCCCGTTA |
| | *M. pulcherrima* | ITS2 | AGCGATATTTACTCTCAAACCTCCGGTTTGGTCCTGCTTCGGCCTAATA<br>TCAACGGCGCTAGAA |
| 511 | *Metschnikowia* sp | D1/D2 | GTACGGCGAGTGAAGCGGCAAAAGCTCCAAATTTGAAATTCCCCCCGGG<br>AATTGTAATTTGAAGAAGATTTGGGTCCGGCCGGCGGGGGTTAAGTCCA<br>CTGGAAAGTGGCGCCACAGAGGGTGACAGCCCCGTGAACCCCTTCAACG<br>CCCTCATCCCAGATCTCCAAGAGTCGAGTTGTTTGGGAATGCAGCTCTA<br>AGTGGGTGGTAAATTCCATCTAAAGCTAAATACCGGCGAGAGACCGATA<br>GCGAACAAGTACAGTGATGGAAAGATGAAAAGCACTTTGAAAAGAGAGT<br>GAAAAAGTACGTGAAATTGTTGAAAGGGAAGGGCTTGCAAGCAGACACT<br>TAACTGGGCCAGCATCGGGGCGGCGGGAAACAAAACCACCGGGGAATGT<br>ACCTTTCGAGGATTATAACCCCGGTCTCAATTTCCTTGCCGCCCCGAGG<br>CCTGCAATCTAAGGATGCTGGCGTAATGGTTGCAAGTCGC |
| | *Metschnikowia* sp | ITS2 | TGATGCGATATTTACTCTCAAACCTCTCGGTTTTCGGTCCTTCGCTTTC<br>GGCCTAATATCAACGGCGCTCGAATAAGTTTTAGCTCATTCTTTTTCCT<br>CACCCTCGTAAGATACCCGCTGAACTTAAGCATACGAATCGAAGGCA |
| 636 | *M. pulcherrima* | D1/D2 | CGGCGAGTGAGCGGCAAAAGCTCAAATTTGAAATCCCCCGGGAATTGTA<br>ATTTGAAGAGATTTGGGTCCGGCCGGCGGGGGTTAAGTCCACTGGAAAG<br>TGGCGCCACAGAGGGTGACAGCCCCGTGAACCCCTTTAACGCCCTCATC<br>CCAGATCTCCAAGAGTCGAGTTGTTTGGGAATGCAGCTCTAAGTGGGTG<br>GTAAATTCCATCTAAAGCTAAATACCGGCGAGAGACCGATAGCGAACAA<br>GTACAGTGATGGAAAGATGAAAAGCACTTTGAAAAGAGAGTGAAAAAGT<br>ACGTGAAATTGTTGAAAGGGAAGGGCTTGCAAGCAGACACTTAACTGGG<br>CCAGCATCGGGGCGGCGGGAAACAAAACCACCGGGGAATGTACCTTTCG<br>AGGATTATAACCCCGGTCCTTACTCCCTTGCTGCCCCGAGGCCTGCAAT<br>CTAAGGATGCTGGCGTAATGGTTGCAAGTCGCCCGT |
| | *M. pulcherrima* | ITS2 | GCGTGGGTGAGCGATATTTACTCTCAAACCTCTGGTTTGGTCCTGCTTC<br>GGCCTAATATCAACGGCGCTGAATAAG |

| | | | |
|---|---|---|---|
| 642 | *M. pulcherrima* | D1/D2 | AAGCTCAAATTTGAAATCCCCCGGGAATTGTAATTTGAAGAGATTTGGG<br>TCCGGCCGGCGGGGGTTAAGTCCACTGGAAAGTGGCGCCACAGAGGGTG<br>ACAGCCCCGTGAACCCCTTTAACGCCCTCATCCCAGATCTCCAAGAGTC<br>GAGTTGTTTGGGAATGCAGCTCTAAGTGGGTGGTAAATTCCATCTAAAG<br>CTAAATACCGGCGAGAGACCGATAGCGAACAAGTACAGTGATGGAAAGA<br>TGAAAAGCACTTTGAAAAGAGAGTGAAAAAGTACGTGAAATTGTTGAAA<br>GGGAAGGGCTTGCAAGCAGACACTTAACTGGGCCAGCATCGGGGCGGCG<br>GGAAACAAAACCACCGGGGAATGTACCTTTCGAGGATTATAACCCCGG |
| | *M. pulcherrima* | ITS2 | CACATTGCGCCCCGGGGTATTCCCCAGGGCATGCGTGGGTGAGCGATAT<br>TTACTCTCAAACCTCCGGTTTGGTCCTGCTTCGGCCTAATATCAACGGC<br>GCTAGAATAAGTTTTAGCCCCAGCCTTTTTCCTCACCCTCGCTAAGAGC<br>TACCTCGCTGTAACTCT |
| 721 | *M. pulcherrima* | D1/D2 | TTTGAAATCCCCCGGGAATTGTAATTTGAAGAGATTTGGGTCCGGCCGG<br>CGGGGGTTAAGTCCACTGGAAAGTGGCGCCACAGAGGGTGACAGCCCCG<br>TGAACCCCTTTAACGCCCTCATCCCAGATCTCCAAGAGTCGAGTTGTTT<br>GGGAATGCAGCTCTAGTGGGTGGTAAATTCCATCTAAAGCTAAATACCG<br>GCGAGAGACCGATAGCGAACAAGTACAGTGATGGAAAGATGAAAAGCAC<br>TTTGAAAAGAGAGTGAAAAAGTACGTGAAATTGTTGAAAGGGAAGGGCT<br>TGCAAGCAGACACTTAACTGGGCCAGCATCGGGGCGGCGGGAAACAAAA<br>CCACCGGGGAATGTACCTTTCGAGGATTATAACCCCGGTCTCAATTTCC<br>TTGTTGCCCCGAGGCCTGCAATCTAAGGATGCTGGCGTAATGGTTGCAA<br>GTCGCCCGTCTG |
| | *M. pulcherrima* | ITS2 | CGTGAATCATTGAATCTTTGAACGCACATTGCGCCCCGGGGTATTCTCT<br>CAGGGCATGCGTGGGTGAGCGATATTTACTCTCAAACCTCCGGTTTGGT<br>CCTGCTTCGGCCTAATATCAACGGCGCTAGAATAAGTTTTAGCCCCAGC<br>CTTTCCTCCTCACCCTCGTAAGACTACCCGCTGAACTATAAGCATA |
| 41 | *M. pulcherrima* | ITS2 | TGAGCGATATTTACTCTCAAACCTCCGGTTTGGTCCTGCTTCGGCCTAA<br>TATCAACGGCGCTAGAATAAGTTTTAGCCCCATCCTTTTTCCTCACCCT<br>CGTAAGACTACCCGCTGAACTTAAGCATATCATAAAGCGGAGGAGAA |
| 99 | *Metschnikowia* sp | ITS2 | GGTGAGCGATATTTACTCTCAAACCTCCGGTTTGGTCCTGCTTCGGCCT<br>AATATCAACGGCGCTAGAATAAGTTTTAGCCCCAGCCTTTTTCCTCACC<br>CTCGTAAGACTACCCGCTGAACTTAAGCATATCATAGAAGCGG |
| 160 | *M. pulcherrima* | ITS2 | TGGGTGAGCGATATTTACTCTCAAACCTCCGGTTTGGTCCTGCTTCGGC<br>CTAATATCAACGGCGCTAGAATAAGTTTTAGCCCCAGCCTTTTTCCTCA<br>CCCTCGTAAGACTACCCGCTGAACTTAAGCATATCA |
| 174 | *M. pulcherrima* | ITS2 | TTTACTCTCAAACCTCCGGTTTGGTCCTGCTTCGGCCTAATATCAACGG<br>CGCTAGAATAAGTTTTAGCCCCATTCTTCTTCCTCACCCTCGTAAGACT<br>ACCCGCTGAACTTAAGCATATCA |
| 186 | *M. pulcherrima* | ITS2 | CATGCGTGGGTGAGCGATATTTACTCTCAAACCTCCGGTTTGGTCCTGC<br>TTCGGCCTAATATCAACGGCGCTAGAATAAGTTTTAGCCCCATTCTTTT<br>TCCTCACCCTCGTAAGACTACCCGCTGAACTTAAGCATATCAGTAAA |
| 204 | *Metschnikowia* sp | ITS2 | AGCGATATTTACTCTCAAACCTCCGGTTTGGTCCTGCTTCGGCCTAATA<br>TCAACGGCGCTAGAATAAGTTTTAGCCCCAGCCTTTTTCCTCACCCTCG<br>TAAGACTACCCGCTGAACTTAAGCATATCATAAAGCGGAGGA |
| 215 | *M. pulcherrima* | ITS2 | TGAATCTTTGAACGCACATTGCGCCCCGGGGTATTCCCCAGGGCATGCG<br>TGGGTGAGCGATATTTACTCTCAAACCTCCGGTTTGGTCCTGCTTCGGC<br>CTAATATCAACGGCGCTAGAATAAGTTTTAGCCCCATTCTTTTTCCTCA<br>CCCTCGTAAGACTACCCGCTGAACTTAAGCATATCA |
| 217 | *M. pulcherrima* | ITS2 | TGAGCGATATTTACTCTCAAACCTCCGGTTTGGTCCTGCTTCGGCCTAA<br>TATCAACGGCGCTAGAATAAGTTTTAGCCCCATTCTTTTTCCTCACCCT<br>CGTAAGACTACCCGCTGAACTTAAGCATATCATAAAGCGGAG |
| 233 | *M. pulcherrima* | ITS2 | GGCATGCGTGGGTGAGCGATATTTACTCTCAAACCTCCGGTTTGGTCCT<br>GCTTCGGCCTAATATCAACGGCGCTAGAATAAGTTTTAGCCCCATTCTT<br>TTTCCTCACCCTCGTAAGACTACCCGCTGAACTTAAGCATATCATAAAG<br>CGGAGG |
| 322 | *M. pulcherrima* | ITS2 | CGCCCCGGGGTATTCCCCAGGGCATGCGTGGGTGAGCGATATTTACTCT<br>CAAACCTCCGGTTTGGTCCTGCTTCGGCCTAATATCAACGGCGCTAGAA<br>TAAGTTTTAGCCCCATCCTTCTTCCTCACCCTCGTAAGACTACCCGCTG<br>AACTTAAGCATATC |

| | | | |
|---|---|---|---|
| 346 | *M. pulcherrima* | ITS2 | ATGCGTGGGTGAGCGATATTTACTCTCAAACCTCCGGTTTGGTCCTGCTTCGGCCTAATATCAACGGCGCTAGAATAAGTTTTAGCCCCATTCTTTTTCCTCACCCTCGTAAGACTACCCGCTGAACTTAAGCATATCATAAAAGCGGAGG |
| 360 | *M. pulcherrima* | ITS2 | GCGATATTTACTCTCAAACCTCTGGTTTGGTCCTGCTTCGGCCTAATATCAACGGCGCTAGAATAAGTTTTAGCCCCAGCCTTTTTCCTCACCCTCGTAAGAGTACCCGCTGAACTTAAGCATATC |
| 380 | *M. pulcherrima* | ITS2 | AGCGATATTTACTCTCAAACCTCCGGTTTGGTCCTGCTTCGGCCTAATATCAACGGCGCTAGAATAAGTTTTAGCCCCATCCTTTTTCCTCACCCTCGTAAGACTACCCGCTGAACTTAAGCATATCAT |
| 564 | *M. pulcherrima* | ITS2 | CATGCGTGGGTGAGCGATATTTACTCTCAAACCTCTGGTTTGGTCCTGCTTCGGCCTAATATCAACGGCGCTAGAATAAGTTTTAGCCCCAGCCTTTTTCCTCACCCTCGTAAGACTACCCGCTGAACTTAAGCATATC |
| 610 | *M. pulcherrima* | ITS2 | TTACTCTCAAACCTCCGGTTTGGTCCTGCTTCGGCCTAATATCAACGGCGCTAGAATAAGTTTTAGCCCCATTCTTTTTCCTCACCCTCGTAAGACTACCCGCTGAACTTAAGCATATCA |
| 623 | *M. pulcherrima* | ITS2 | CCCCAGGGCATGCGTGGGTGAGCGATATTTACTCTCAAACCTCCGGTTTGGTCCTGCTTCGGCCTAATATCAACGGCGCTAGAATAAGTTTTAGCCCCATCCTTTTTCCTCACCCTCGTAAGACTACCCGCTGAACTTAAGCATATCATAAAGCGGAGGA |
| 660 | *M. pulcherrima* | ITS2 | CCCCAGGGCATGCGTGGGTGAGCGATATTTACTCTCAAACCTCCGGTTTGGTCCTGCTTCGGCCTAATATCAACGGCGCTAGAATAAGTTTTAGCCCCAGCCTTTTTCCTCACCCTCGTAAGACTACCCGCTGAACTTAAGCATATA |
| 669 | *M. pulcherrima* | ITS2 | GGGTGAGCGATATTTACTCTCAAACCTCCGGTTTGGTCCTGCTTCGGCCTAATATCAACGGCGCTAGAATAAGTTTTAGCCCCATCCTTTTTCCTCACCCTCGTAAGACTACCCGCTGAACTTAAGCATAC |
| 692 | *M. pulcherrima* | ITS2 | AGCGATATTTACTCTCAAACCTCCGGTTTGGTCCTGCTTCGGCCTAATATCAACGGCGCTAGAATAAGTTTTAGCCCCATCCTTTTTCCTCACCCTCGTAAGACTACCCGCTGAACTTAAGCATATCAT |
| 702 | *M. pulcherrima* | ITS2 | AGCGATATTTACTCTCAAACCTCCGGTTTGGTCCTGCTTCGGCCTAATATCAACGGCGCTAGAATAAGTTTTAGCCCCAGTCTTTTTCCTCACCCTCGTAAGACTACCCGCTGAACTTAAGCATATCAT |

**Supplementary Table 6.** Pul1–4 primary sequences from genome–available *K. lactis, C. auris* and *Metschnikowia* species reported to produce iron chelators and Pul1/2 *Metschnikowia* isolates from this study. The CTG codons from yeasts that belong to the CTG–clade have been translated to serine instead of leucine. Loci with two ORFs have been included as *a* and *b*.

| **Pul1** | |
|---|---|
| yAMV240 | MVTESGELFISQAFIDMWIDSLVGYLPGETCRHVSQFVQNEYVGTLGQLYVTIRDL<br>RQAISAFVDSVNGEENKNFLAFDAHVSDCGCHLRASMSMDLIQRYRGNREELLSFL<br>GLVDACDNALVSTSALMKDICTEAKSLKELQLPKSTKDPLLFLNAIGWKFESDNLS<br>EIKYIFYCYVLSQFKTYSFRNK<br>QDSVHIDTDKEFKQKNEMICTHTCQGKGKLGNGCRYLKHARIGKAALKQWTLCYQE<br>RLSKMSVDYLAKSDSELKELVENSRKESHKSVAAVPSYVQFKISERLWAFNQFPFL<br>LSMRVFVDENHEDIYARAFVGRDLKWNIQFVTSDVLEDTPHIIVAGHCRVPHGYND<br>TNKLNLANLSLDAHQNMRSFWYSFMSQHKQYPFDTALGCDDDLQNVLPAHEFKDYM<br>KFKYAGIRAFRDMEFTPKHIFVEYPSVVFSKQRMLAGKQGVLFI |
| yAMV312 | MVTESGELFISQAFIDMWIDSLVGYLPGETCRHVSQFVQNEYVGTLGQLYVTIRDL<br>RQAISAFVDSVNGEENKNFLAFDAHVSDCGCHLRASMSMDLIQRYRGNREELLSFL<br>GLVEACDKALVSTSVLMKDICTEAKSLKELQLPKSTKDPLLFLNAIGWKFESNNLS<br>EIKYIFYCYVLSQFKTYSFRNKQDSVHIDTDKEFKQKNEMICTHTCQGKGKLGNGC<br>RYLKHARIGKAALKQWTLCYQERLSKMSVVYLAKSDSELKELVENSRKESHKSVAA<br>VPSYVQFKISERLWASNQFPFLLSMRVFVDEGHEDIYARAFVGRDLKWNIQFVTSD<br>VLEDTPHIIVAGHCRVPHGYNDINTLNLANLSLDAHQNMRSFWYSFMSQHKQYPFD<br>TALGCDDDLQNVLPAHEFKDYMKFKYAGIRAFKDMEFTPKHIFVEYPSVVFSKQRM<br>LAGKQGVLFI |
| yAMV460 | MVTESGELFISQAFIDMWIDSLVGYLPGETCRHVSQFVQNEYVGTLGQLYVTIRDL<br>RQAISAFVDSVNGEENKNFLAFDAHVSDCGCHLRASMSMDLIQRYRGNREELLSFL<br>GLVEACDNALVSTSALMKDICTEAKSLKELQLPKSTKDPLLFLNAIGWKFESNNLS<br>EIKYIFYCYVLSQFKTYSFRNKQDSVHIDTDKEFKQKNEMICTHTCQGKGKLGNGC<br>RYLKHARIGKAALKQWTLCYQERLSKMSVDFLAKSDSELKELVENSRKESHKSVAA<br>VPSYVQFKISERLWASNQFPFLLSMRVFVDEGHEDIYARAFVGRDLKWNIQFVTSD<br>VLEDTPHIIVAGHCRVPHGYNDTNKLNLANLSLDAHQNMRSFWYSFMSQHKQYPFD<br>TALGCDDDLQNVLPAHEFKDYMKFKYAGIRAFRDMEFTPKHIFVEYPSVVFSKQRM<br>LAGKQGVLFI |
| yAMV511 | MVTESGELFISQAFIDMWIDSLVGYLPGETCRHVSQFVQNEYVGTLGQLYVTIRDL<br>RQAISAFVDSVNGEENKNFLAFDAHVSDCGCHLRASMSMDLIQRYRGNREELLSFL<br>GLVEACDRALVSTSALMKDICTEAKSLKELQLPKSTKDPLLFLNAIGWKFESNNLS<br>EIKYIFYCYVLSQFKTYSFRNKQDSVHIDTDKEFKQKNEMICTHTCQGKGKLGNGC<br>RYLKHARIGKAALKQWTLCYQERLSKMSVDYLAKSDSELKELVETSRKESHKSVAA<br>VPSYVQFKISERLWASNQFPFLLSMRVFVDEGHDDIYARAFVGRDLKWNIQFVTSD<br>VLEDTPHIIVAGHCRVPHGYNDINKLDLANLSLDAHQNMRSFWYSFMSQHKQYPFD<br>TALGCDDDLQNVLPAHEFKDYMKFKYAGIRAFRDMEFTPKHIFVEYPSIVFSKQRM<br>LAGKQGVLFI |
| yAMV636 | MVTESGELFISQAFIDMWIDSLVGYLPGETCRHVSQFVQNEYVGTLGQLYVTIRDL<br>RQAISAFVDSVNGEENKNFLAFDAHVSDCGCHLRASMSMDLIQRYRGNREELLSFL<br>GLVDACDNALVSTSALMKDICTEAKSLKELQLPKSTKDPLLFLSAIGWKFESNNLS<br>EIKYIFYCYVLSQFKTYSFRNKQDSVHIDTDKEFKQKNEMICTHTCQGKGKLGNGC<br>RYLKHARIGKAALKQWTLCYQERLSKMSVDYLAKSDSELKELVETSRKESHKSVAA<br>VPSYVQFKISERLWAFNQFPFLLSMRVFVDEGHEDIYARAFVGRDLKWNIQFVTSD<br>VLEDTPHIIVAGHCRVPHGYNDTNKLNLANLSLDAHQNMRSFWYSFMSQHKQYPFD<br>TALGCDDDLQNVLPAHEFKDYMKFKYAGIRAFRDMEFTPKHIFVEYPSIVFSKQRM<br>LAGKQGVLFI |
| yAMV642 | MVTESGELFISQAFIDMWIDSLVGYLPGETCRHVSQFVQNEYVGTLGQLYVTIRDL<br>RQAISAFVDSVNGEENKNFLAFDAHVSDCGCHLRASMSMDLIQRYRGNREELLSFL<br>GLVEACDNALVSTSALMKDICTEAKSLKELQLPKSTKDPLLFLNAIGWKFESNNLS<br>EIKYIFYCYVLSQFKTYSFRNKQDSVHIDTDKEFKQKNEMICTHTCQGKGKLGNGC |

| | |
|---|---|
| | RYLKHARIGKAALKQWTLCYQERLSKMSVDYLAKSDSELKELVENSRKESHKSVAA<br>VPSYVQFKISERLWASNQFPFLLSMRVFVDEGHEDIYARAFVGRDLKWNIQFVTSD<br>VLEDTPHIIVAGHCRVPHGYNDTNKLNLANLSLDAHQNMRSFWYSFMSQHKQYPFD<br>TALGCDDDLQNVLPAHEFKDYMKFKYAGIRAFRDMEFTPKHIFVEYPSVVFSKQRM<br>LAGKQGVLFI |
| *M. pulcherrima* APC1.2 | MVTESGELFISQAFIDMWIDSLVGYLPGETCRHVSQFVQNEYVGTLGQLYVTIRDL<br>RQAISAFVDSVNGEENKNFLAFDAHVSDCGCHLRASMSMDLIQRYRGNRKELLSFL<br>GLVDACDNALVSTSALMKDICTEAKSLKELQLPKSTKDPLLFLNAIGWKFESNNLS<br>EIKYIFYCYVLSQFKTYSFRNKQDSVHIDTDKEFKQKNEMICTHTCQGKGKLGNGC<br>RYLKHARIGKAALKQWTLCYQERLSKMSVDYLAKSDSELKELVENSRKESHKSVAA<br>VPSYVQFKISERLWAFNQFPFLLSMRVFVDEGHEDIYARAFVGRDLKWNIQFVASD<br>VLEDTPHIIVAGHCRVPHGYNDTNKLNLANLSLDAHQNMRSFWYSFMSQHKQYPFD<br>TALGCDDDLQNVLPAHEFKDYMKFKSAGIRAFKDMEFTPKHIFVEYPSVVFSKQRM<br>LAGKQGVLFI |
| *M. pulcherrima* AP47 | MVTESGELFISQAFIDMWIDSLVGYLPGETCRHVSQFVQNEYVGTLGQLYVTIRDL<br>RQAISAFVDSVNGEENKNFLAFDAHVSDCGCHLRASMSMDLIQRYRGNREELLSFL<br>GLVEACDKALVSTSVLMKDICTEAKSLKELQLPKSTKDPLLFLNAIGWKFESNNLS<br>EIKYIFYCYVLSQFKTYSFRNKQDSVHIDTDKEFKQKNEMICTHTCQGKGKLGNGC<br>RYLKHARIGKAALKQWTLCYQERLSKMSVDYLAKSDSELKELVENSRKVSHKSVAA<br>VPSYVQFKISERLWASNQFPFLLSMRVFVDESHEDIYARAFVGRDLKWNIQFVTSD<br>VLEDTPHIIVAGHCRVPHGYNDTNKLNLANLSLDAHQNMRSFWYSFMSQHKQYPFD<br>TALGCDDDLQTVLPAHEFKDYMKFKYAGIRAFRDMEFTPKHIFVEYPSVVFSKQRM<br>LAGKQGVLFI |
| *M. pulcherrima* KIOM G15050 (*a*) | MVTESGELFISQAFIDMWIDSLVGYLPGETCRHVSQFVQNEYVGTLGQLYVTIRDL<br>RQAISAFVDSVNGEENKNFLAFDAHVSDCGCHLRASMSMDLIQRYRGNREEILSFL<br>GLVEACDKALVSTSALMKDICTEAKSLKELQLPKSTKDPLLFLNAIGWEI |
| *M. pulcherrima* KIOM G15050 (*b*) | MHPHLPRKGKLGNGCRYLKHARIGKAALKQWTLCYQERLSKMSVDYLAKSDSELKE<br>LVENSRKESHKSVAAVPSYVQFKISERLWASNQFPFLLSMRVFVDEGHKDIYARAF<br>VGRDLKWNIKFVTSDVLEDTPHIIVAGHCRVPHGYNDINKFNLANLSLDAHQNMRS<br>FWYSFMSQHKQYPFDTALGCDDDLQNVLPAHEFKDYMKFKYAGIRAFRDMEFTPKH<br>IFVEYPSVVFSKQRMLAGKQGVLFI |
| *M. rubicola* CBS 15344 | MVTESGELFISQAFIDMWIDSLVGYLPGETCRHLSQFVQNEYVGTLGQLYVTIRDL<br>RSAISAFVDSVNGEENKNFLAFDAHVSDCGCHLRASMSMDLIQRYRGNREEILSFL<br>GLVEACDKALVSTSALMKDICTEAKSLKELQLPKSTKDPLLFLNAIGWKFESDNLS<br>EIKYIFYCYVLSQFKTYSFRNKQDSVHIDTDKEFKQKNEMICTHTCQGKGKLGNGC<br>RYLKHARIGKAALKQWTLCYQERLSKMSVDYLAKSDSELKELVENSRKESHKSVAA<br>VPSYVQFKISERLWAFNQFPFLLSMRVFVDEGHEDIYARAFVGRDLKWNIQFVTSD<br>VLEDTPHIIVAGHCRVPHGYNDINKLNLANLSLDAHQNMRSFWYSFMSQHKQYPFD<br>TALGCDDDLQNVLPAHEFKDYMKFKYAGIRAFRDMEFTPKHIFVEYPSIVFSKQRM<br>LAGKQGVLFI |
| *M. chrysoperlae* NRRL Y-27615 | MVTESGELFISHPFTDMWIDSLLGYLPGETCRHVSQFVQNEYVGTLGQLYVTIMDL<br>RQAVSAFVDSVNGEENKNFLAFDAHVSDCGCHLRASMSMDLIQRYRKNKKELILFR<br>GLVDACDNALVSTSALMKDICTEAKSLKELQLPKSTKDPLLFLNAIGWKFESKNLS<br>EIKYIFYCYVLSQFKTYSFRNKQDSVHIDTDMEFKQKNEMICTHTCQGKGKLGNGC<br>RYLKHARIGKAALKQWTLCYQERLSKMSVDYLAKCDSELRELVENSRKESHKSVAA<br>VPSYVQFKISERLWASNEFPFLLSMRVFVDEGHEDIYARAFVGRDLRWEIRSVTNQ<br>ELKDTPHIVVAGHCHVPQGCDDISKLDLETFSLDMHQNMRSFWYSFMSQHKQYPFD<br>TPLGCDDELQDVLPAHEFEDYMKFKYTGIRAFGDIEFTPKHIFVEYPSVVFNKQRM<br>LAGKQGVLFI |
| *C. auris* B11221 | MWIDTLLGLLPGDKYFQQTFKRNEYVGALAQLCVTIMDLKLASEAFLGSIEGKETF<br>MPFDAHVSDCGCHLRAQLSMEKVFHYKKHPEKLEVFRKVIEVCEGSLLRASELSKA<br>LCWSEQSPLEFGLPKQTATRDDFKIAIEWVSDKFDGNSDIKYVYYCYTLSRFKRYN<br>ERNLKDSVEFDLDAASKCLNSLVYGFTCQGKGKLGEGCRYLKHAKIGKGFVKTQTT<br>ALQARLSQMSVEYLVSLVPELSNELKTYCVKSKKGIYAVPALLQYKVLERVWSARR<br>TPLFLAIRFFDGDLSKFYYTFAMVAKDLQWAFHNESKRNEPHIAITFDCEISGLAC |

| | |
|---|---|
| | LKDDDLSNVMCNLQPKMRETWYTFMTQHKQYPFELCGSLEQDLEAKIHSEVEVPKF<br>RNTFHEPNGANLEPKHIFLDYPRAVTEKQASISGKAGTLLIN |
| *K. lactis* NRRL Y-1140 | MYQLLFQRLGVTLTAGNDKKTSIPSNQLVGHLIGLILLCDDLNEAFADFQALLQNG<br>IAISSSDRGYLVFDAHVTDCGCHLRAQMIQDVFSYFKNHEVTKLYIFDAVAKKLNE<br>LKRHCISLIQTLCWENTSAQKLGFPKNIKSYEELLKLLQWDTADISSAIADSFPMN<br>PNNADQNEKGLVWNVTKLEVVLEFLYCSHFLSKNKIYNKKENLDSVTIDFNNAFEK<br>RQLLSNKFHCQGTGKLGVGCRYLKHAKQSKNSFISVVSNLQSRLALLSIAFLKSRC<br>TLPCDIEALQKNSPRNVSAIPNFLHFLILEREWSQNETPILLAVRKLHEHEHCDLY<br>FEARINPHTFEWTLQHKECCEFEHHTPYIVITALATGSSTTKTAQLLAWELMKAQK<br>NFRQFWLTFMSQHRQYPFEIEHDEDQLLETQVSQDIFELYCQSKREDRNQILFDDS<br>TSLLPKHIFTEYPSIFFNFQKNVCSKHGALVI |
| **Pul2** | |
| yAMV240 | MLTILSFPVFWVLVVSLCLVSSKSKLRAFQKQPPFKKIPRSHRNVEGKKITQGPEI<br>SEKNKNQYGSIYSHRDGFRYEVVLTTPSQLKQYYSSHTKDHKKLDSFGAGQYLVAL<br>LGECLGFQNGESWTRMRKSFNLFFTHTLAAKTLPAMIAFIDGWISEHDSQNEFSVD<br>AFDFVATVPFTCIAKYLYGSEQCSGRVLTELKNLVPLHSELMTHAFTTFWGRFKIY<br>QYFPFQRMKDLKFFQDSFKSLSLAMVESARDAENPTVASELYRLVESRDLTLDNWI<br>QSLDEILFANIDVTATIMSWSLVEMGRNKHEQARLRLEVLENLHSMDEYCKRTDTV<br>LHRVLLEILRLHPLLWYGFPEQSSSAMVIDGHKIEANTPIVVDQYQLNYKSPLWNP<br>TDKSTDYGATFDSNRFLGLNNRDILMSSVTFGSGPRRCLGKNFAEVLIKTEIAKVL<br>STFEVALEGELKVAADTFVVRPDAQIKLTRLI |
| yAMV312 | MITILSFPVFWVLVVSLCLVSSKSKLRAFQKQPPFKKIPRSHRNVEGKKITQGPEI<br>SEKNKNQYGSIYSHRDGFRYEVVLTTPSQLKQYYSSHTKDHKKLDSFGAGQYLVAL<br>LGECLGFQNGESWTRMRKSFNLFFTHTLAAKTLPAMIAFIDGWISEHDSQDEFSVD<br>AFDFVATVPFTCIAKYLYGNEQCSGRVLTELKNLVPLHSELMTHAFTTFWGRFKIY<br>QYFPFQRMKDLKFFQDSFKSLSLAMVESARDAENLTVASELYKLVESRDLTLDNWI<br>QSLDEILFANIDVTATIMSWSLVEMGRNKHEQARLRLEVLENLQSVDEYCKRTDTV<br>LHRVLLEILRLHPLLWYGFPEQSSSAMVIDGHKIEANTPIVVDQYQLNYKSPLWNP<br>ADKSTDYGATFDSDRFLGLNNRDILMSSVTFGSGPRRCLGKNFAEVLIKTEIAKVL<br>STFEVALEGELKVAADTFVVRPDAQIKLTRLI |
| yAMV460 | MITILSLPVFWVLVVSLCLVSSKSKLRAFQKQPPFKKIPRSHRNVEGKKITQGPEI<br>SEKNKNQYGSIYSHRDGFRYEVVLTTPSQLKQYYSSHTKDHKKLDSFGAGQYLVAL<br>LGECLGFQNGESWTRMRKSFNLFFTHTLAAKTLPAMIAFIDGWISEHDSQNEFSVD<br>AFDFVATVPFTCIAKYLYGNEQCSGRILTELKNLVPLHSELMTHAFTTFWGRFKIY<br>QYFPFQRMKDLKFFQDSFKSLSLAMVESARDAENPTVASELYKLVESRDLTLDNWI<br>QSLDEILFANIDVTATIMSWSLVEMGRNKHEQARLRLEVLENLQSVDEYCKRTDTV<br>LHRVLLEILRLHPLLWYGFPEQSSSAMVIDGHKIEANTPIVVDQYQLNYKSPLWNP<br>ADKRTDYGATFDSDRFLGLNNRDILMSSVTFGSGPRRCLGKNFAEVLIKTEIAKVL<br>FTFEVALEGELKVAADTFVVRPDAQIKLTRLI |
| yAMV511 | MLTILSLPVFWVLVVSLCLVSSKSKLRAFQKQPPFKKIPRSHRNVEGKKITQGPEI<br>SEKNKNQYGSIYSHRDGFRYEVVLTTPSQLKQYYSSHTKDHKKLDSFGAGQYLVAL<br>LGECLGFQNGESWTRMRKSFNLFFTHTLAAKTLPAMIAFIDGWISEHDSQNEFSVD<br>AFDFVATVPFTCIAKYLYGNEQCSGRVLTELKNLVPLHSELMTHAFTTFWGRFRIY<br>QYFPFQRMKDLKYFQDSFKSLSLAMVESARDAENPTVASELYKLVESRDLTLDNWI<br>QSLDEILFANIDVTATIMSWSLVEMGRNKHEQARLRLEVLENLQSVDEYCKRTDTV<br>LHRVLLEILRLHPLLWYGFPEQSSSAMVIDGHKIEANTPIVVDQYQLNYKSPLWNP<br>ADKSTDYGATFDSNRFLGLNNRDILMSSVTFGSGPRRCLGKNFAEVLIKTEVAKVL<br>STFEVALEGELKVAADTFVVRPDAQIKLTRLI |
| yAMV636 | MLTILSLPVFWVLVVSLCLVSSKSKLRAFQKQPPFKKIPRSHRNVEGKKITQGPEI<br>SEKNKNQYGSIYSHRDGFRYEVVLTTPSQLKQYYSSHTKDHKKLDSFGAGQYLVAL<br>LGECLGFQNGESWTRMRKSFNLFFTHTLAAKTLPAMIAFIDGWISEHDSQDEFSVD<br>AFDFVATVPFTCIAKYLYGSEQCSGQVLTELKNLVPLHSELMTHAFTTFWGRFKIY<br>QYFPFQRMKDLKFFQDSFKSLSLAMVESARDAENPTVASELYKLVESRDLTLDNWI<br>QSLDEILFANIDVTATIMSWSLVEMGRNKHEQARLRLEVLENLQSVDEYCKRTDTV<br>LHRVLLEILRLHPLLWYGFPEQSSSAMVIDGHKIEANTPIVVDQYQLNYKSPLWNP |

| | |
|---|---|
| | ADKSTDYGATFDSDRFLGLNNRDILMSSVTFGSGPRRCLGKNFAEVLIKTEIAKVL<br>STFEVALEGELKVAADTFVVRPDAQIKLTRLI |
| yAMV642 | MLTILSLPVFWVLVVSLCLVSSKSKLRAFQKQPPFKKIPRSHRNVEGKKITQGPEV<br>SEKNKNQYGSIYSHRDGFRYEVVLTTPSQLKQYYSSHTKDHKKLDSFGAGQYLVAL<br>LGECLGFQNGESWTRMRKSFNLFFTHTLAAKTLPAMIAFIDGWISEHDSQNEFSVD<br>AFDFVATVPFTCIAKYLYGNEQCSGRVLTELKNLVPLHSELMTHAFTTFWGRFKLY<br>QYFPFQRMKDLKFFQDSFKSLSLAMVESARDAENPTVASELYKLVESRDLTLDNWI<br>QSLDEILFANIDVTATIMSWSLVEMGRNKHEQARLRLEVLENSQSVDEYCKRTDTV<br>LHRVLLEILRLHPLLWYGFPEQSSSAMVIDGHKIEANTPIVVDQYQLNYKSPLWNP<br>ADKSTDYGATFDSDRFLGLNNRDILMSSVTFGSGPRRCLGKNFAEVLIKTEIAKVL<br>STFEVALEGELKVAADTFVVRPDAQIKLTRLI |
| *M. pulcherrima*<br>APC1.2 | MITILSFPVFWVLVVSLCLVSSKSKLRAFQKQPPFKKIPRSHRNVEGKKITQGPEI<br>SEKNKNQYGSIYSHRDGFRYEVVLTTPSQLKQYYSSHTKDHKKLDSFGAGQYLVAL<br>LGECLGFQNGESWTRMRKSFNLFFTHTLAAKTLPAMIAFIDGWISEHDSQNEFSVD<br>AFDFVATVPFTCIAKYLYGNEQCSGRILTELKNLVPLHSELMTHAFTTFWGRFKIY<br>QYFPFQRMKDLKFFQDSFKSLSLAMVESARDAENPTVASELYKLVESRDLTLDNWI<br>QSLDEILFANIDVTATIMSWSLVEMGRNKHEQARLRLEVLENLQSVDEYCKRTDTV<br>LHRVLLEILRLHPLLWYGFPEQSSSAMVIDGHKIEANTPIVVDQYQLNYKSPLWNP<br>ADKRTDYGATFDSDRFLGLNNRDILMSSVTFGSGPRRCLGKNFAEVLIKTEIAKVL<br>FTFEVALEGELKVAADTFVVRPDAQIKLTRLI |
| *M. pulcherrima*<br>AP47 (*a*) | MLTILTFPVFWVLVVSLCLVSSKSKLRAFQKQPPFKKIPRSHRNVEGKKITQGPEI<br>SEKNKNQYGSIYSHRDGFRYEVVLTTPSQLKQYYSSHTKDHKKLDSFGAGQYLVAL<br>LGSVWAFKMVSCGLGCGNCLICFSHIRWSRKRCPR |
| *M. pulcherrima*<br>AP47 (*b*) | MRKLFNLFFTHTLEPKTLPAMIAFIDGWISEHDSQNEFSVDAFDFVATVPFTCIAK<br>YLYGNEQCSGRVLTELKNLVPLHSELMTHAFTTFWGRFKLYQYFPFQRMKDLKFFQ<br>DSFKSLSLAMVESARDAENPTVASELYKLVESRDLTLDNWIQSLDEILFANIDVTA<br>TIMSWSLVEMGRNKQEQGRLRLEVLENLQSVDEYCKRTDTVLHRVLLEILRLHPLL<br>WYGFPEQSSSAMVIDGHKIEANTPIVVDQYQLNYKSPLWNPADKSTDYGATFDSDR<br>FLGLNNRDILMSSVTFGSGPRRCLGKNFAEVLIKTEVAKVLSTFEVALEGELKVAA<br>DTFVVRPDAQIKLTRLI |
| *M. pulcherrima*<br>KIOM G15050 | MITILSLPVFWVLVVSLCLVSSKSKLRAFQKQPPFKKIPRSHRNVEGKKITQGPEI<br>SEKNKNQYGSMYSHRDGFRYEVVLTTPSQLKQYYSSHTKDHKKLDSFGAGQYLVAL<br>LGECLGFQNGESWTRMRKSFNLFFTHTLAAKTLPAMIAFIDGWISEHDSQNEFSVD<br>AFDFVATVPFTCIAKYLYGNEQCSGQVLTELKNLVPLHSELMTHAFTTFWGRFKIY<br>QYFPFQRMKDLKFFQDSFKSLSLAMVESARDAENLTVASELYKLVESRDLTLDNWI<br>QSLDEILFANIDVTATIMSWSLVEMGRNKHEQARLRLEVLENLQSVDEYCKRTDTV<br>LHRVLLEILRLHPLLWYGFPEQSSSAMVIDGHKIEANTPIVVDQYQLNYKSPLWNP<br>ADKSTDYGATFDSDRFLGLNNRDILMSSVTFGSGPRRCLGKNFAEVLIKTEIAKVL<br>STFEVALEGELKVAADTFVVRPDAQIKLTRLI |
| *M. rubicola*<br>CBS 15344 | MITILSFPVFWVLVVSLCLVSSKSQLRAFQKQPPFKKIPRSHRNVEGKKITQGPEI<br>SEKNKNQYGSIYSHRDGFRYEVVLTTPSQLKQYYSSHTKDHKKLDSFGAGQYLVAL<br>LGECLGFQNGESWTRMRKSFNLFFTHTLAAKTLPAMIAFIDGWISEHDSQDEFSVD<br>AFDFVATVPFTCIAKYLYGNEQCSGRVLTELKNLVPLHSELMTHAFTTFWGRFKIY<br>QYFPFQRMKDLKFFQDSFKSLSLAMVESARDAENPTVASELYKLVESRDLTLDNWI<br>QSLDEILFANIDVTATIMSWSLVEMGRNKHEQARLRLEVLENLQSVDEYCKRSDTV<br>LHRVLLEILRLHPLLWYGFPEQSSSAMVIDGHKIEANTPIVVDQYQLNYKSPLWNP<br>ADKSTDYGATFDSDRFLGLNNRDILMSSVTFGSGPRRCLGKNFAEVLIKTEIAKVL<br>STFEVALEGELKVAADTFVVRPDAQIKLTRLI |
| *M. chrysoperlae*<br>NRRL Y-27615 | MITVLSFPVFWVLVVSLCLVTSKSKLRAFQKQPPFKKIPRSHRNVEGKKITEGPEI<br>SEKNKKQYGSIYSHRDGFRYEVVLTTPSQLKQYYSSHTKDHKKLDSFGAGQYLVAL<br>LGECLGFQNGESWTRMRNSFNLFFTHTLAAKTLPAMIAFIDGWISEHDSQNEFSVD<br>AFDFVATVPFTCIAKYLYGDEQCSGRVLTELKNLVPLHSELMTHAFTTFWGRFKIY<br>QYFPFQRMKDLKLFQDSFKSLSLAMVESARDAENPTVASELYKLVESRDLTLDNWI<br>QSLDEILFANIDVTATIMSWSLVEMGRNRHEQVRLRLEVLENLQSVDEYCKRTDTV<br>LHRVLLEILRLHPLLWYGFPEQSSSDMVIDGHKIEANTPIVVDQYQLNYKSPLWNP |

| | |
|---|---|
| | ADKSTDYGATFDSDRFLGLNNRDILMSSVTFGSGPRRCLGKNFAEVLIKTEIAKVL<br>STFEVALEGELKVAADTFVVRPDAQIKLKRLI |
| *C. auris* B11221 | MSNLHVDLILVLVVATIFFSRKIIDQPNFSKLLKGSRNIPYTNRNVARYKLSKGSE<br>ISVENKSKLGTLYMHRDGFRYEVVLTTPDQLKQYFSSHPRDHAKLDSFGAGQYLVA<br>LLGECLGFQNGDSWMNMRKLFNGYFSHASAIQTLPTMVEFIGGYVESTTKESSKVV<br>DPFEYVSAIPFTCIANYLYGKSHCTPERLSGLKELIPLHTDLLTFAFKTFWGRFKI<br>FQFFQSDEMKGLRHFQDRFTQLSLEMVRNAGNDPSVASELYSRVEQGELSFDSWIQ<br>TLDEILFANIDVTATVMSWALVEMARNPNAQHAVREEIHSSTESQEEYVKKTETHL<br>HRVLLETLRLHPLLWFGFPEVVSKDIVIDGFQIPAHTPIVIDQFQVNYESEIWNPP<br>NKPKGFGHEFHPDRFIGLGNRDVLMSSVTFGSGPRRCLGKNFAEIVVKTLLVNMVK<br>HFELSLCDPVEYAENTFVVQPKTKVKLGRIGV |
| *K. lactis* NRRL Y-1140 | MLADILIPLIKKNWMAFVYFTPVLFVVLYLLKEWRAAYGFNNLGQTVAAPFGYERK<br>TLPYNKENCARTKFLDGKSLSIKNRDQCGDLYLQRSGTYKEIVLTTPKQLMEYYKS<br>NSKNHSKLDSFGAGAFLVALLGECLGFQNGSEWLSMRKVFDSFFTHKAAVENFPVM<br>IDYISEWIKDLDTEQISDIDPLQLVSDLPFTCIAKYLYGSELCSKQFLQALKDLIP<br>MHTELMHYSFLTVAGRFKIFQYFPSKKMKQVSQFQRQFIDLSLKQVELSRQSGQET<br>VVEKLYRHVESGKFTFNNWIQTIDEILFANIEVTSTVMAWALVEMGSNIEEQNRLR<br>CEILKVKEQSSKDDFNKETDPMQRYMKLTDTYLQYCVWETLRMHPLLWFSFPEISS<br>ETLFIDGIRISPNTPIVVDQYQINYNSPIWNPSDKPKDFGKKFAPSRFENITLRDA<br>LYSQVTFGAGSRKCLGRNFAELLIKSELAYILSKYKVTLTEKVEFSKDTFVVQPKT<br>KIQLTAL |
| **Pul3** | |
| yAMV511 | MAISQLVAIFASYCDFKIHNFKVNSNNGPTFVCSFIILAVTALLVFVLENPAVPTK<br>KTSMSFTKALQGFFSAPKWTLAGGIILLWGMFFASFLMSEVVYFMPVFLTESLGWE<br>TKFQGVAFMVASIIGICGSLVFPHLIEMPIRKKEKEFNSQAESQSKLSEKSDISHN<br>SEESADRKQEFEAYKKNALYNNQIVLSLVSLLIALVGQAFMIGAAQVFRHRSLPHI<br>NTGCFFVGGMSLVMLAYNGMASTFPALFSVYIDPQVKLQLMPAIGAVAALGKLIAP<br>IVLSNLYQTKLGLSIAVGLGMILTALTLMPVFLLKDKKH |
| *M. pulcherrima* APC1.2 | MAISQLVAIFASYCDFKIHNFKVNSNNGPTFVCSFIILAVTALLVFVLENPAVPTK<br>KTSLSFTKALQGFFSAPKWTLAGGIILLWGMFFASFLMSEVVYFMPVFLTESLGWE<br>TKFQGVAFMVASIIGICGSLVFPHLIEMPIRKKEKELNSQAESQSKLSEKSDISHN<br>SEESADRKQEFEAYKKNALYNNQIVLSLVSLLIALVGQAFMIGAAQVFRHRSLPHI<br>NTGCFFVGGMSLVMLAYNGMASTFPALFSVYIDPQVKLQLMPAIGAVAALGKLIAP<br>IVLSNLYQTKLGLSIAVGLGMILTALTLMPVFLLKDKKH |
| *M. pulcherrima* AP47 | MAISQLVAIFASYCDFKIHNFKVNSNNGPTFVCSFIILAVTALLVFVLENPAVPTK<br>KTSMSFTKALQGFFSAPKWTLAGGIILLWGMFFASFLMSEVVYFMPVFLTESLGWE<br>TKFQGVAFMVASIIGICGSLVFPHLIEMPIRKKEKEFNSQAESQSKLSEKSDISHN<br>SEESADRKQEFEAYKKKALYNNQIVLSLVSLLIALMGQAFMIGAAQVFRHRSLPHI<br>NTGCFFVGGMSLVMLAYNGMASTFPALFSVYIDPQVKLQLMPAIGAVAALGKLIAP<br>IVLSNLYQTKLGLSIAVGLGMILTALTLMPVFLLKDKKH |
| *M. pulcherrima* KIOM G15050 | MAISQLVAIFASYCDFKIHNFKVNSNNGPTFVCSFIILAVTALLVFVLENPAVPTR<br>KTRMSFTKALQGFFSAPKWTLAGGMILLWGMFFASFLMSEVVYFMPVFLTESLGWE<br>TKFQGVAFMIASIIGICGSLVFPHLIEMPIRKKEKEFNSQAESQSKLSEKSDISHN<br>SEESADRKQEFEAYKKNALYNNQIVLSLVSLLIALVGQAFMIGAAQVFRHRSLPHI<br>NTGCFFVGGMSLVMLAYNGMASTFPALFSVYIDPQVKLQLMPAIGAVAALGKLIAP<br>IVLSNLYQTKLGLSIAVGLGMILTALTLMPVFLLKDKKH |
| *M. rubicola* CBS 15344 | MAISQLVAIFASYCDFKIHNFKVNSNNGPTFVCSFIILAVTALLVFVLENPAVPTK<br>KTSMSFTKALQGFFSAPKWTLGGGIILLWGMFFASFLMSEVVYFMPVFLTESLGWE<br>TKFQGVAFMVASIIGICGSLVFPHLIEMPIRKKEKEFNSQAESQSRLSEKSDISHN<br>SEESADRKQEFEAYKKNALYNNQIVLSLVSLLIALVGQAFMIGAAQVFRHRSLPHI<br>NTGCFFVGGMSLVMLAYNGMASTFPALFSVYIDPQVKLQLMPAIGAVAALGKLIAP<br>IVLSNLYQTKLGLSIAVGLGMILTALTLMPVFLLKDKKH |
| *M. chrysoperlae* NRRL Y-27615 | MAISQLVAIFASYCDFKIHNFKVNSNNGPTFVCSFIILAVTALLVFVLENPAVPTK<br>KTSMSFTKALQGFFTAPKWTLAGGIILLWGMFFASFLMSEVVYFMPVFLTESLGWE<br>TKFQGVAFMVASIIGICGSLVFPHLIEMPIRKKEKELNAQAESQSKLSEKSDTSQN |

| | |
|---|---|
| | SEDSADRKQEFEAYKKNALYNNQVVLSLVSLLIALVGQAFMIGASQVFRHRSLPHI<br>NTGCFFVGGMSLVMLAYNGMASTFPALFSVYIDPQVKLQLMPAIGAVAALGKLIAP<br>IVLSNLYQTKLGLSIAVGLGMILTALTLMPVFLLKIKKH |
| *C. auris* B11221 | MALSQLVAIFGSYCDFLIGDFRVSSNNAPTFITSFIMVLVAMGLFFTMKNPPVPKR<br>KNTISFKRALLDFFTARRAQLLGSVIIMWGMFLASFLMSEVVYFMPVFLTQSLGWE<br>TKFQGVAFMVASLIGIIGSLLCPKLLSKLTENKEPEIHFESEKASTDESLEEMRKK<br>QTEEFKKRQVFRNQVVLTLVSLAFALVGQAFMIGANEIFNDRRLPHINVGCFFVAG<br>MSIIMLGYNGMASTFPSMFSEYVKPEVKVQLMPIIGAMTALGKLVAPIVLANLYTT<br>RLGLSIAVALGIILTGITVPPTYCLIIS |
| *K. lactis* NRRL Y-1140 | MKLTDSQKHLYSQYLAVTLIAVQFSFDTCVYLSSVVQYVKECGSDDPENYLFILQA<br>VSAAVQVFFSFIIGDIASYVGSIKWVIIFLYFLSFVGNFLYSCAGAVSLNTLLGGR<br>IICGAASSSGAVVYSYITAISKDRTTIFKLFSIYRTSAGICMALAQLVAILFALCD<br>FTVRGYRITSYNAPTFASSFIILLICVLLMFVLENPPVKSARNPKNYLDAWKKFFS<br>AGSNRLIASLILLWNMFLSTFFMCEVLYFMPIFLTLNVGWKTEYEGVAFMVSAVLG<br>VAGSFFAPDLVKLFAKLNTPSTQDETDTSDNDKIEKEESEQKSDINTLHRNQVSLT<br>IFALFVALIGQAFMIGASEALSNDKLPKTNSGIFFTAGLSITMLGYNFMGSSVPAL<br>FSMYIDPQVKVQLMPFIGAIAGVGKLVAPIVLAALYKTPLGLPIGVGFGMILVGIS<br>IPSLVYLRRNKM |
| *S. cerevisiae* 288C | MSIAQDRGIVFKLLSIYRAAAGIFMALAQLIVIFFGYCDFKIKGYRIASYNAPTFA<br>SSFIILAVCLLLVVVLENPEVKVTNSENSLFSALKQFFRVERKKLISCLILLWSMF<br>LSSFIMSEVVYFMPLFLTLHVNWDTKFQGIAFMVASILGVTGSYFAPKLINVGCSC<br>GRAKDGGLEESDTTGSETVEVKKKDSLYSGQVFLSIFALFVSLLGQAFMIGASEAL<br>KHKSMPPTNSGIFFSAGMSITLLGYNFLASSIPALFSMYIDPKLKVQLMPSIGAIS<br>GIGKLVAPIVLAALYGTRLGLSIAVGFGMILVAVSIPPLIWLRKKRC |
| **Pul4** | |
| *M. pulcherrima* APC1.2 | MVKVAKTCTRCKSKKLKCDGSKPSCSRCLSLGITSCEYPPDKRVDKRQKLNGKDVF<br>VFKNCDTTGDYKDIPETTLISPISQIEAGLDGSLSTFNASEFPTFFPSPDIGQQWP<br>DFVPDLFQGDLNFSMDEFFCFLDDSVPLGSYDPLEQIEISPHKLLIDAVFSNKLHP<br>PPGVSHEMILNIDMLQVRAEDEEFLHATILAMGALTLAKRDLVGQRQDTMVEVPEK<br>IGLKLPAMASEAYGHYSRARELIPSMLKAPSRHGFRGLAVMANFMSILLTPETQMY<br>ISYHALQIAISIGLTQMSRVEINTEEYGLVIAFWGLWCSSCMLATFHGHAPPLMRS<br>KISTTAHVQMKNRYVEVFFELRIKFAELSGIVAVVNKSEEINYQLLRKELSRLCEE<br>ISLFEESFPQEEIIGRHELFTLELKCWKAYVSMLLNLPALLEKRNTRAVVDAKNIV<br>KDFWAYYYFYKRNFLPNLDWNFSYPLRNATLCVWFACMVLTRYINSSPALSFEFAE<br>YKIGMELLIDLTGVIPINKYLIKELENVKNNELDSDKWLFKQG |
| *M. pulcherrima* AP47 | MVKVAKTCTRCKSKKLKCDGSKPSCSRCLSLGITSCEYPADKRVDKRQKLNGKDVF<br>VFKNCDTTGDYKDIPETTLISPISQIEAGLDGSLSTFNASEFPTFFPSPDIGQQWP<br>DFVPDLFQGDLNFSMDEFFCFLDDSVPLGSYDPLEQIEISPHKLLIDAVFSNKLHP<br>PPGVSHEMILNIDMLQVRAEDEEFLHATILAMGALTLAKRDLVGQRQDTMVEVPEK<br>VGLKLPAMASEAYGHYSRARELIPSMLKAPSRNGFRGLAVMANFMSILLTPETQMY<br>ISYHALQIAISIGLTQMSRVELDTEEYGLVIAFWGLWCSSCMLATFHGHAPPLMRS<br>KISTTAHVQMKNRYVEVFFELRIKFAELSGIVAVVNKSEEINYQLLRKELSRLCEE<br>ISLFEESFPQEEIIGRHELFTLELKCWKAYVSMLLNLPALLEKRNSRAVVDAKNIV<br>KDFWAYYYFYKRNFLPNLDWNFSYPLRNATLCVWFACMVLTRYINSSPALSFEFAE<br>YKIGMELLTDLTGVIPINKYLIKELENVKNNELDSDKWLFKQG |
| *M. pulcherrima* KIOM G15050 (*a*) | MVKVAKTCTRCKLKKTQM |
| *M. pulcherrima* KIOM G15050 (*b*) | MASEAYGHYSRARELIPSMLKAPSRHGFRGLAVMANFMSILLTPETQMYISYHALQ<br>IAISIGLTQMSRVEIDTEEYGLVIAFWGLWCSSCMLATFHGHAPPLMRSKISTTAH<br>VQMKNRYVEVFFELRIKFAELSGIVAVVNKSEEINYQLLRKELSRLCEEISLFEES<br>FPQEEIIGRHELFTLELKCWKAYVSMLLNLPALLEKRNSRAVVDAKNIVKDFWAYY<br>YFYKRNFLPNLDWNFSYPLRNATLCVWFACMVLTRYINSSPALSFEFAEYKIGMEL<br>LIDLTGVIPINKYLIKELENVKNNELDSDKWLFKQG |

| | |
|---|---|
| *M. rubicola* CBS 15344 | MVKVAKTCTRCKSKKLKCDGSKPSCSRCLSLGITSCEYPPDKRVDKRQKLNGKDVF<br>VFKNSDTTGDHKDVPETTLISPISQIEAGLDASLSTFSATEFPTFFPSPDIGQQWP<br>DFVPDLFQGDLNFSMDEFFCFLDDSVPLGSYDPLEQVEIPPPKLLLDAVFSNKLHP<br>PPGVSYEMILNIDMLQVRAEDEEFLHATILAMGALTLAKRDLVGQRQDTMVEVPEK<br>VGLKLPAMASEAYGHYSTARELIPSMLKAPSRHGFRGLAVMANFMSILLTPETQMY<br>ISYHALQIAISIGLTQVSRVEIDTEEYGLIIAFWGLWCSSCMLATFHGHAPPLMRS<br>KITTTAHVQMKNRYVEIFFELRIKFAELSGIVAVVNKSEEINYQLLRKELSRLCEE<br>ISLFEESFPQEYIIGRHELFTLELKCWKAYVSMLLNLPALLEKRNSRAVVDAKNIV<br>KDFWAYYYFYKRNFLPNLDWNFSYPLRNATLCVWFACMVLTTYINSSPALSFEFAE<br>YKIGMELLTDLTGVIPINKYLIKELENVKNNQLEVDKWLFKQE |
| *M. chrysoperlae* NRRL Y-27615 | MVKVAKTCTRCKSKKLKCDGSKPSCSRCLSLGITSCEYPPDKRVDKRQKLNGKDVF<br>VFKNSDTTGDHKDVPETTLISPISQIEAGLDASLSTFSATEFPTFFPSPDIGQQWP<br>DFVPDLFQGDLNFSMDEFFCFLDDSVPLGSYDPLEQVEIPPPKLLLDAVFSNKLHP<br>PPGVSYEMILNIDMLQVRAEDEEFLHATILAMGALTLAKRDLVGQRQDTMVEVPEK<br>VGLKLPAMASEAYGHYSTARELIPSMLKAPSRHGFRGLAVMANFMSILLTPETQMY<br>ISYHALQIAISIGLTQVSRVEIDTEEYGLIIAFWGLWCSSCMLATFHGHAPPLMRS<br>KITTTAHVQMKNRYVEIFFELRIKFAELSGIVAVVNKSEEINYQLLRKELSRLCEE<br>ISLFEESFPQEYIIGRHELFTLELKCWKAYVSMLLNLPALLEKRNSRAVVDAKNIV<br>KDFWAYYYFYKRNFLPNLDWNFSYPLRNATLCVWFACMVLTTYINSSPALSFEFAE<br>YKIGMELLTDLTGVIPINKYLIKELENVKNNQLEVDKWLFKQE |
| *C. auris* B11221 | MARACVTCKHKKLKCDGERPACGRCCKLGLPCEIPRDKRSDKRKVENGTSSFVFKN<br>TGRKGRKHGWDKPVAIDDNELSSAPPLEASLGGEFVRGSLLEPTTLAWPELDEVKL<br>PYDLDIEEFLHSLDYPLGLETTEPVGKVTSVFTGFSPVGEDQPTPYHVLIDTVFSD<br>ASHTPVSITKAMITGIADKVDRSEDESFLLSIVLAIGALTLAKQKYAEQRQSILKL<br>GTDASDPSYHIKTPLVANEAVNHYQEARKLNSHILENPSAHGFRGLVLISNFLSGL<br>LTLEAQMFISFHALQVAVAIKMHTNKETGSSEEESGLKIAFWELWCSACLFASFHG<br>RLPPIRREEITTSLDFDLKNEYNKRFFHLRVEIAELHCQVAALKVNGLASSVSERD<br>FHTKVSTISQRIVELEALFPQTESNFYRQELLTLELKCWMSQAIMLEGQQRLTRKL<br>SIKSVVEARKLIRELWSYYNPEKFARGTMLSHLDWNFTYPLRTATIGAFTAAKTLT<br>LFINSVDYLTFDFFDYCLSRKVLEFLTGVMSINKHLLQDLDNYALR |
| *K. lactis* NRRL Y-1140 | MACLECKKRKQKCDGQKPCRRCTKLNVKCIYGTDRRKDKRKIKDGSNMFIFKNQTL<br>CNDKINGIVPHPLSHDTITTKETWEPSYPLFSDDINPADIISMENTDGSIPLQFDL<br>DFTSLESCDVNDFLRLIGDTFPANDADTLDMNQMGGFNTPSISHNTQDDKSQIAIQ<br>RNRLIDVIFGDDSHTPPGILREHIFELSERHEDLEALDFDDNGKFLLSTVLCLGAL<br>TLRKRELLNRDSNQPSTNGIPEVAAGAYKYYTIATDLIPAVHAAPNIDGFCGLVLM<br>ANFMTIMIPLEGQLYLSNNALEVAVALNFHKRESYDEMIVSNPAQLGVFLLFWNLW<br>CSSCMLATLLGKQPFLTLDNISLPPPHQMQHTVSSSPLSINFMRLRIQLATLQTKI<br>FQRLYVYGSLNKVLFQEIETELSLLSTQISNMKCYPIYDEGLFYRSKVLMLELSCL<br>KAHNAFLLYRPNLIQKKSLHAVDAAKHIILEIWSHYTKQFPKNEKDLVDHLDWNFS<br>YPLRTASLTLSISCVILQKYQQSLNFLEEYGIFEYNLALGVLNDLIQVVPIEKRLI<br>NLLTVSRTTVEDANESNREDSLRFWTNMLMC |
| *S. cerevisiae* 288C | MDRSKDARKRSISLACTVCRKRKLKCDGNKPCGRCIRLNTPKECIYNIDKRKDKRK<br>IKNGSKVFLFKNNTIDNGNNSILENKGLNEDLSSHIYEKEAPKFDSDIDISRFGTN<br>DAVIFNNDGWDTSLPIDFDFDEFNTETTDFDDFLKLLGDNSPSKEQKSLSYSPTAT<br>GLSGVVKETESEDNAPTRSRLIDVLFENKLHSVPGISKWHLYELESQYPNLECTEG<br>NSDEKFLLSTVLCLGSLTIRKRELLNHSNIDNRPLLPENSISKLTTDAFKYYNAAK<br>TLVPDLLSHPTIDGFCGLVLMANFMTMMISLEHQLYLSINALQLAVALNLNNNTKC<br>KELLESNSDGIGVILLFWNIWCSSCMLATIHGKNPFITLEQITTPLPCEISPRNKT<br>NKLLIDFMQIRIKLATLQSKIFQRLYTSSTANEVPFVNLEREFEEVSLQITRLKGF<br>PIFEEHLFYRSRVLMLELSCLRAQASFLLYRPYLITGESLQAVTMAKSIIHEIWSQ<br>YTKQFPDNEKERHERLDWNFCYPLRTASLTLCISCIILLRYKQVVQFLKGTELFEY<br>ILALEILQDLVQVLPIEQNLIDIIKYPISPVQLSGDSFVEFWGRILY |

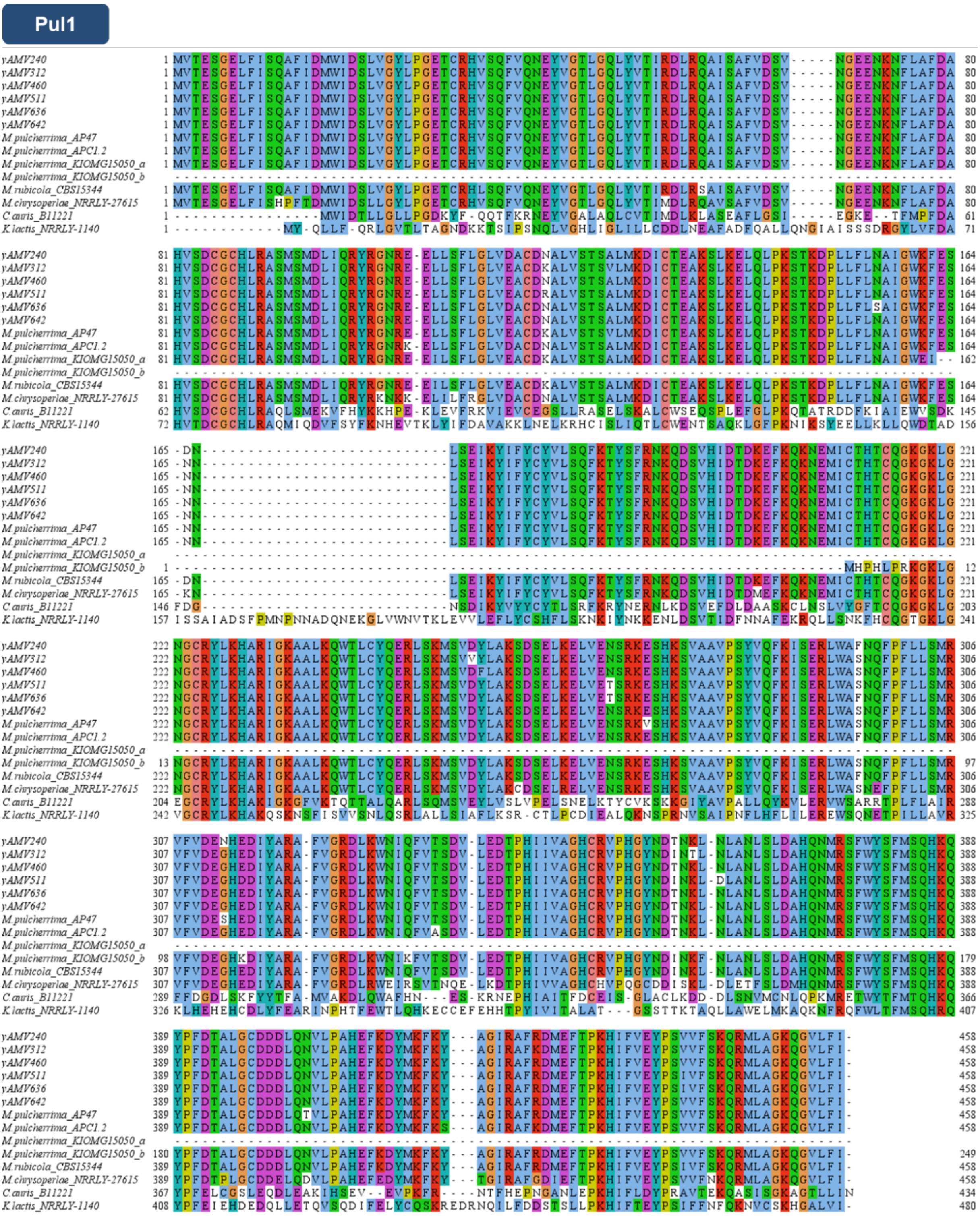


**Supplementary Figure 1**. Multiple sequence alignment of Pul1 from *Metschnikowia* species, *C. auris* and *K. lactis* from NCBI databank and yeast isolates from this study. Amino acids are highlighted using Clustal Colour Scheme. All *Metschnikowia* strains show high similarity but a unique Pul1 sequence. For *M. pulcherrima* KIOM G15050, two ORFs were identified that aligned to different parts of the Pul1 proteins of other organisms, shown as *a* and *b*.

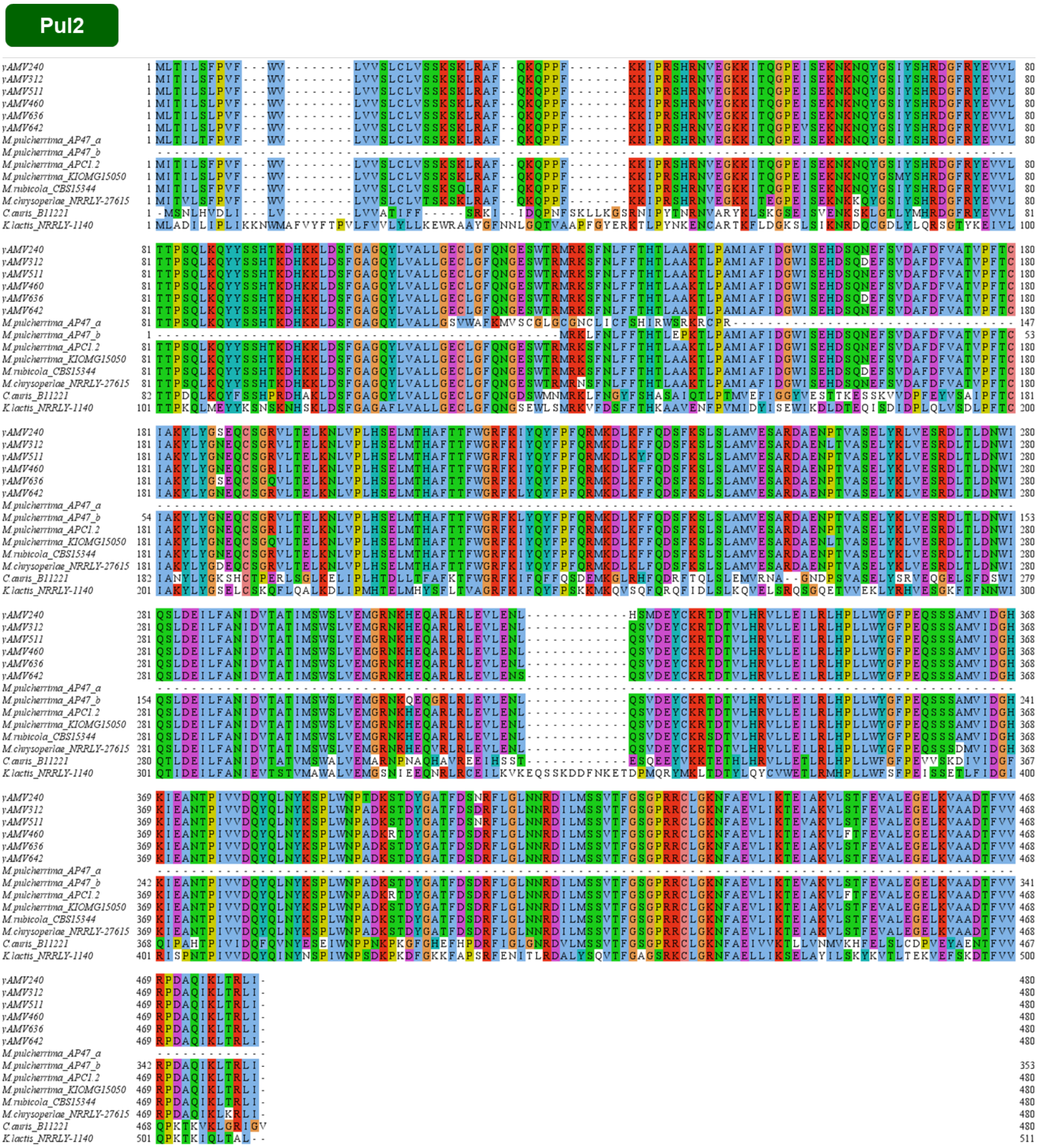


**Supplementary Figure 2.** Multiple sequence alignment of Pul2 from *Metschnikowia* species, *K. lactis* and *C. auris* from NCBI databank and yeast isolates from this study. Amino acids are highlighted using Clustal Colour Scheme. All *Metschnikowia* strains show high similarity but a unique Pul2 primary protein sequence. For *M. pulcherrima* AP47, two ORFs were identified that aligned to different parts of the Pul2 proteins of other organisms, shown as *a* and *b*.

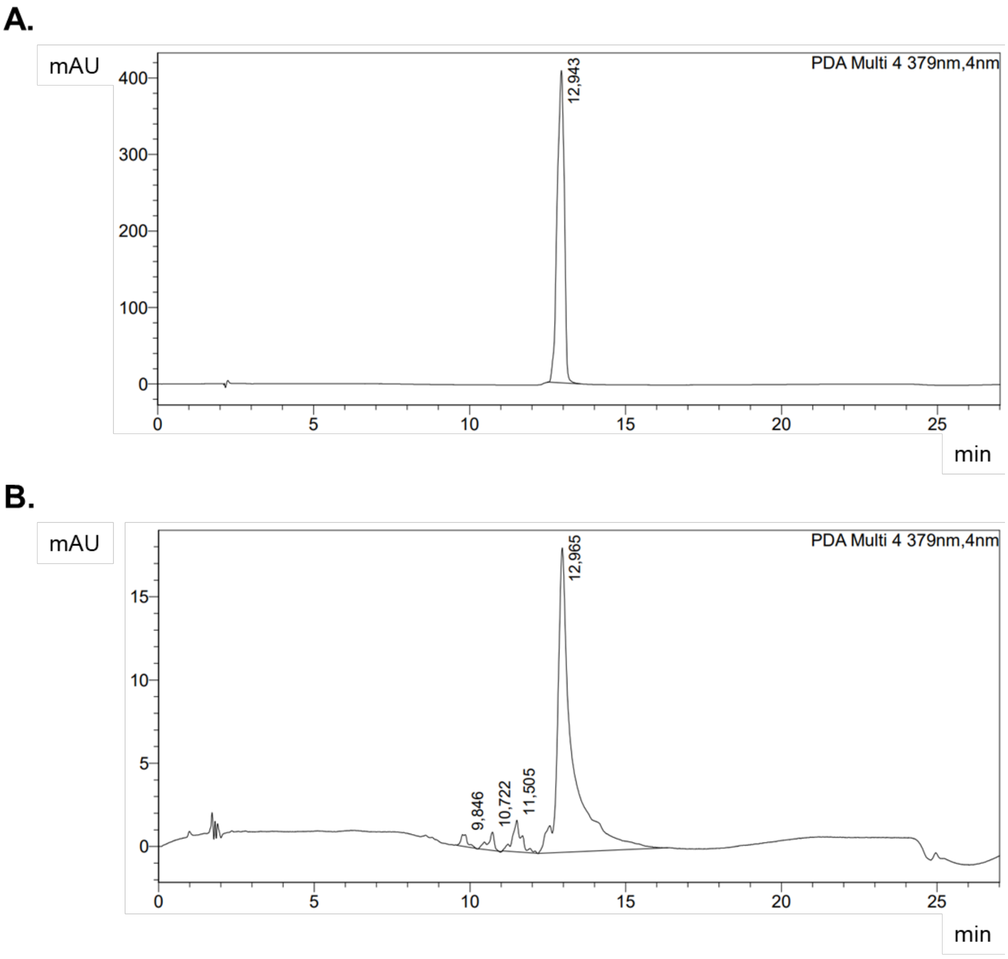


**Supplementary Figure 3**. HPLC chromatogram of (**A**) pulcherriminic acid standard dissolved in DMSO and (**B**) 50–fold concentrated spent media from yAMV511 isolate culture at 379 nm. Both samples show a pick at the same retention time.

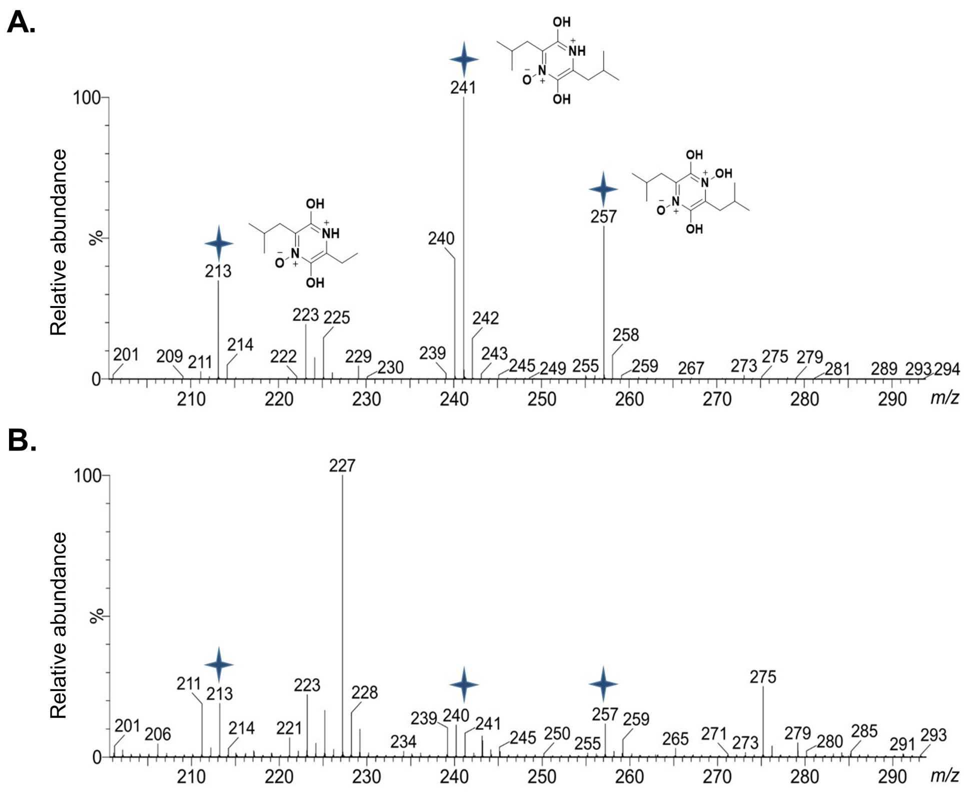


**Supplementary Figure 4.** Mass spectrometry profile of (**A**) pulcherriminic acid standard dissolved in DMSO and (**B**) 50–fold concentrated spent media from yAMV511 isolate culture. The peaks with highest intensity in the standard (highlighted and with the chemical structure) can be found in the spent media of yAMV511 as well, indicating pulcherriminic acid production.

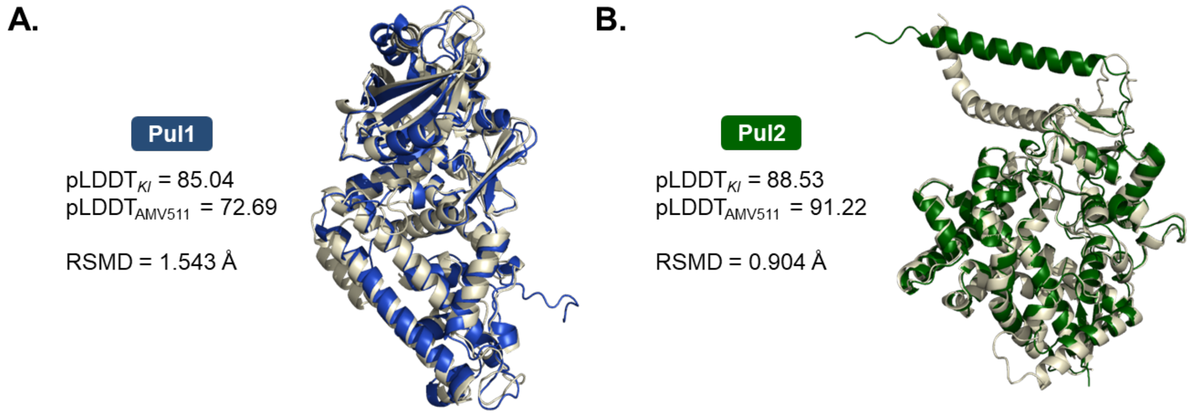


**Supplementary Figure 5.** Pul2 shows a more similar structure than Pul1 when comparing Pul1/2 from *K. lactis* and yAMV511 isolate. **A.** Alignment of AlphaFold–predicted structures of Pul1 for yAMV511 (dark blue) and *K. lactis* (beige). **B.** Alignment of AlphaFold–predicted structures of Pul2 for yAMV511 (dark green) and *K. lactis* (beige).
